# Mapping Endogenous Psychedelic Chemistry: Indolethylamine *N*-Methyltransferase (INMT) Localizes Proximal to the Golgi Apparatus of Human Neurons

**DOI:** 10.64898/2026.10.30.758666

**Authors:** Karis Will, Sergio Lázaro Martínez, Surya A. Reis, Stephen J. Haggarty

## Abstract

*N,N*-dimethyltryptamine (DMT) is a serotonergic psychedelic with a history of ethnobotanical use and emerging clinical efficacy for neuropsychiatric disorders. Following Julius Axelrod’s pioneering studies showing DMT production in human tissues, DMT has been proposed as an endogenous neuromodulator because it is a serotonin 2A receptor (5-HT_2A_ R) agonist. However, the DMT-producing enzyme, indolethylamine *N*-methyltransferase (INMT), has not been characterized at the protein level in the human central nervous system. Here, we validated an anti-INMT antibody and used immunocytochemistry to show that INMT localizes in close proximity to other DMT biosynthetic enzymes near the cytosolic face of the *cis*-Golgi network in human induced pluripotent stem cell-derived neurons, with immunoreactivity observed across multiple neuronal Golgi phenotypes. Notably, both DMT-like and 5-HT_2A_ R immunoreactivity also partially localized in close proximity to the Golgi apparatus. Our results establish the first protein-level characterization of INMT in human neurons and identify the *cis*-Golgi network as a candidate microdomain for psychedelic tryptamine biosynthesis. We propose that INMT catalytic activity at the *cis*-Golgi network creates an enriched region of membrane-permeable psychedelic tryptamines that can access an intracellular 5-HT_2A_ R pool. These findings provide new tools and insights that can inform efforts to understand the mechanism of action of psychedelics under clinical investigation and to target endogenous tryptamines therapeutically.

**GRAPHICAL ABSTRACT:** 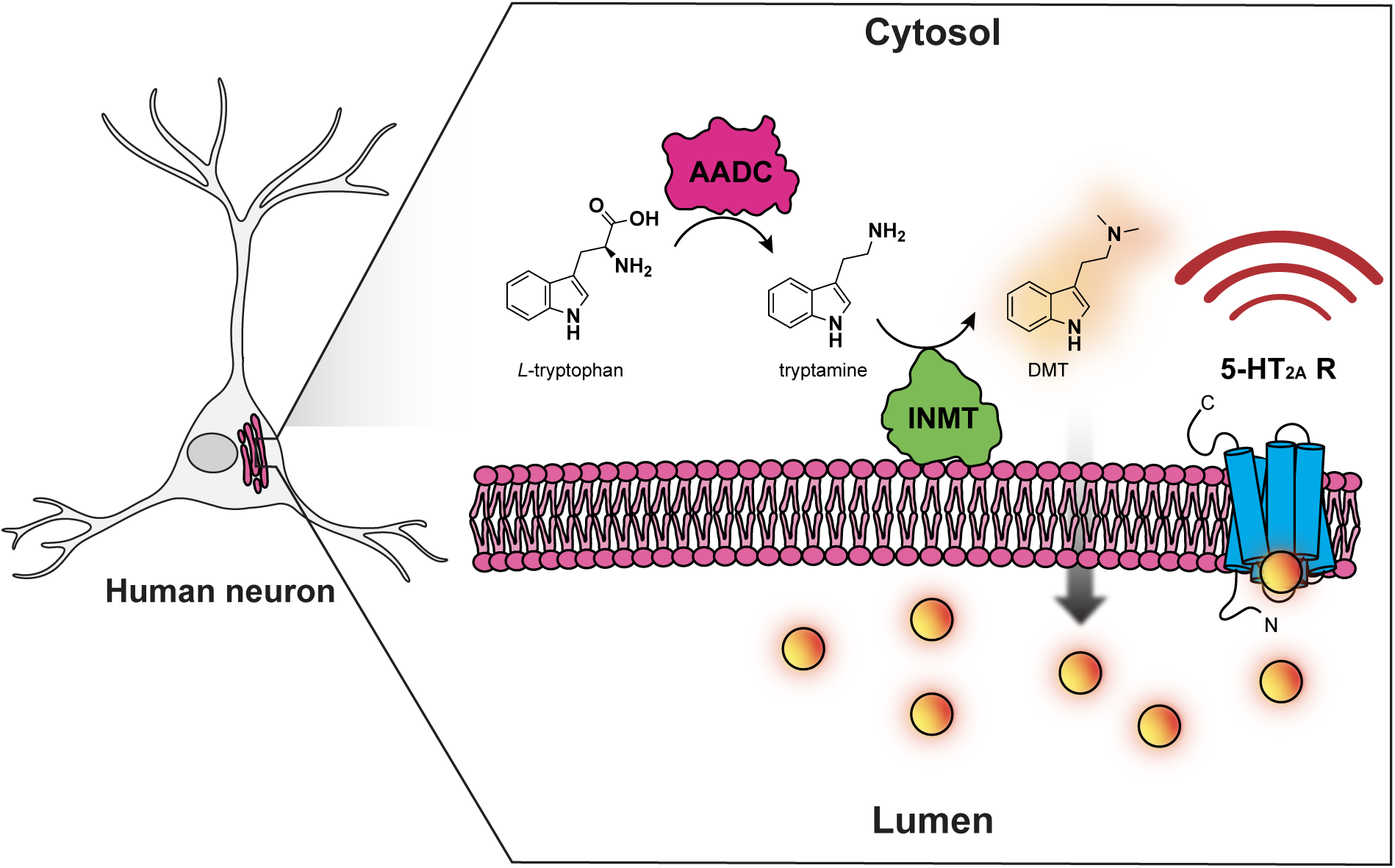

## INTRODUCTION

Indolethylamine *N-*methyltransferase (INMT; EC 2.1.1.49) is a transmethylation enzyme that facilitates the *N*-methylation of various small molecule amine substrates. INMT is the candidate enzyme responsible for synthesizing the classic psychedelic compound *N,N-*dimethyltryptamine (DMT) in mammals.^1,2,3^ DMT is commonly characterized as a secondary plant metabolite and principal psychoactive component of ayahuasca, a traditional medicine used in many Indigenous cultures of South America.^4^ When administered exogenously, DMT and related serotonergic psychedelics, such as psilocybin and lysergic acid diethylamide (LSD), induce profound alterations to mood, perception, and cognition in humans via agonism of the serotonin 2A receptor (5-HT_2A_ R).^5^ In traditional contexts, practitioners use DMT to achieve ancestral divination and treat physical and mental ailments.^6^ In the clinic, exogenously administered serotonergic psychedelics induce rapid and sustained antidepressant effects even after a single dose, in contrast to conventional antidepressants that often require weeks of use before symptom alleviation.^7,8,9,10,11,12,13,14^ A hallmark of major depressive disorder is a loss of dendritic spine density in the prefrontal cortex, which is hypothesized to lead to cortical dysregulation.^15^ At the cellular level, DMT promotes structural plasticity in cortical neurons, including increased dendritic complexity and spine density, suggesting that its observed therapeutic effects in humans are linked to restored neuronal structure.^16^ Despite DMT’s extensive ethnobotanical use and growing clinical interest, it remains unclear whether the human central nervous system (CNS) synthesizes DMT endogenously. Furthermore, the cellular and subcellular environments in which INMT-mediated DMT biosynthesis may occur are underexplored, despite the growing recognition of the importance of spatial organization in cellular signaling processes.

Emerging evidence challenges the historical focus on G protein-coupled receptor (GPCR) signaling at the plasma membrane, and introduces the concept of location bias, which posits that receptor activation can result in distinct signaling outcomes dependent on its subcellular location.^17,18,19^ A recent report highlights this phenomenon in rat cortical neurons, demonstrating that a 5-HT_2A_ R population localizes to the Golgi apparatus, and activating intracellular 5-HT_2A_ Rs contributes to downstream changes in structural plasticity.^20^ The authors indicate that intracellular 5-HT_2A_ R signaling is distinct from plasma membrane signaling and may contribute to the neuroplasticity-inducing properties of psychedelics. In neurons, the Golgi apparatus is a critical hub for protein trafficking and plays a key role in cellular polarization, positioning it as a functionally relevant site for intracellular signaling cascades.^21^ Localizing INMT, the enzyme that methylates tryptamine and serotonin, near or at the Golgi apparatus would clarify the role of endogenous psychedelic tryptamines and intracellular 5-HT_2A_ Rs.

The ability of mammalian tissues to biosynthesize methylated tryptamines is well established, originating from landmark studies in the 1960s by Axelrod and colleagues.^22^ These early efforts led to the identification of an *N*-methyltransferase, INMT, in mammalian brain homogenates that converts supplied serotonin, *N*-methylserotonin, tryptamine, and *N*-methyltryptamine (NMT) to *N*-methylserotonin, bufotenine, NMT, and DMT, respectively.^22^ Based on these studies and cross-comparison to the biosynthesis of known indoleamine neurotransmitters, the putative mammalian DMT biosynthetic pathway was established (**Scheme 1**).^23,24^ Like indoleamine neurotransmitter biosynthesis, DMT production starts with the essential amino acid L-tryptophan. L-tryptophan is first decarboxylated and converted to tryptamine by the ubiquitously expressed cytosolic enzyme, aromatic L-amino acid decarboxylase (AADC; 4.1.1.28). Tryptamine is subsequently methylated via INMT in a two-step process at the primary aliphatic amine, first yielding NMT, followed by DMT. Notably, serotonin is also an established substrate of INMT, which is methylated at the analogous primary amine to generate bufotenine, another methylated tryptamine with documented traditional use and emerging clinical efficacy.^22,8,25^ The catabolism of DMT and NMT is primarily regulated by monoamine oxidase A (MAO-A; EC 1.4.3.4), a flavin adenine dinucleotide-dependent oxidative deaminase.

**Scheme 1.**
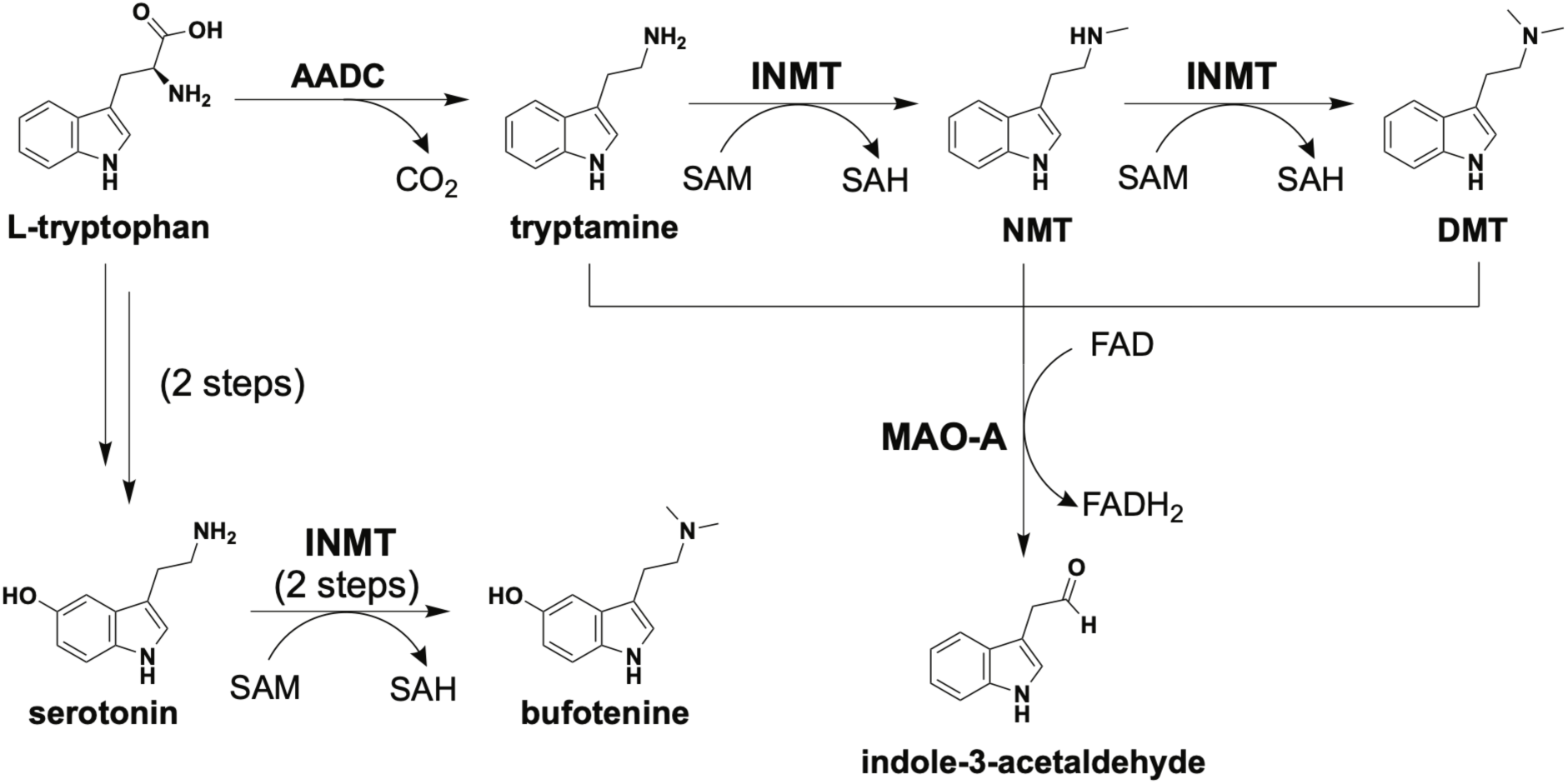
Putative mammalian methylated tryptamine biosynthetic pathway.

Human, rabbit, mouse, and rat brain homogenates have all shown *N*-methylation of tryptamine or NMT activity *in vitro*, yet detecting endogenously synthesized DMT *in vivo* remains an analytical challenge (**Table 1**).^26,27^ Recent reports have demonstrated for the first time the detection and quantification of endogenous DMT and bufotenine in rodent brain, where their concentrations were all within the low to subnanomolar range and comparable to serotonin (**Table 1**).^28,29,30^ Given that global CNS concentrations of DMT and related indolamines are often below the limit of detection of traditional mass spectrometry systems, researchers commonly use INMT as a proxy for DMT production, analogous to the use of tryptophan hydroxylase as a marker for serotonin biosynthesis.^31^ *INMT* messenger ribonucleic acid (mRNA) has to date been characterized in human, mouse, and rat brain using single-cell detection techniques like single-cell RNA sequencing (scRNA-seq) and *in-situ* hybridization (ISH) (**Table 1**).^28,32,33,34^ Interestingly, in a bulk detection method, rabbit brain tissue yielded very low to no detection of *INMT* mRNA.^35^ This low abundance of *INMT* transcripts in a bulk analysis method suggests that INMT expression may be localized to specific brain nuclei or subcellular domains in the CNS.^36^ This hypothesis was supported in an ISH experiment revealing that *INMT* mRNA is expressed in discrete cell types and regions of postmortem human brain (**Table 1**).^28^ Despite robust characterization of *INMT* mRNA in mammalian brain, detection of INMT at the protein level and whether it is spatially positioned to support DMT biosynthesis remains unknown (**Table 1**). Mapping INMT at the protein level in the human brain is critical to uncovering DMT’s endogenous role as mRNA alone does not reflect protein abundance or localization. Furthermore, elucidating INMT’s subcellular localization in human neurons can provide insight into its enzymatic activity and potential neurobiological function.

**Table 1:** Evidence for DMT production and INMT detection in mammalian brain tissues and cell types. Table assembled from data found in references 26 - 35. No data reflects a lack of evidence in the literature.

| **Species** | **DMT Synthesis *in vitro*** | **Endogenous DMT** | ***INMT* mRNA Expression** | **INMT Protein Expression** |
|---|---|---|---|---|
| Human | Demonstrated | No data | Detected<br>(scRNA-seq, ISH) | No data |
| Rabbit | Demonstrated | No data | Low abundance<br>(Northern blot) | No data |
| Mouse | Demonstrated | Detected | Detected<br>(scRNA-seq, ISH) | No data |
| Rat | Demonstrated | Detected | Detected<br>(ISH) | No data |

Human INMT (hINMT) is a 263 amino acid *S*-adenosyl-L-methionine (SAM)-dependent methyltransferase with a well-defined catalytic site and resolved crystal structure (PDB: 2A14). A pairwise sequence alignment analysis of mammalian INMTs revealed that human and rabbit INMTs share ∼88% amino acid sequence identity, but rodent INMTs show lower similarity with human and rabbit INMTs (**Figure 1a**). Consistent with this, phylogenetic tree analysis revealed that primate and rabbit INMTs form a clade separate from rodent and ungulate INMTs (**Figure 1b**). To test whether these divergences contribute to structural differences that affect substrate binding, we performed *in silico* docking analyses to compare tryptamine binding at the catalytic site of human and rodent INMTs. This effort identified a binding site variant, L201F, present in rodent INMTs that potentially introduces steric clash with the tryptamine substrate and alters the size of the substrate binding pocket (**Figure 1c**). These genetic and structural differences may contribute to species-specific substrate tolerability and could explain why DMT production is preserved in INMT knockout (KO) rodent models, and rat INMT catalytic inactivity towards tryptamine.^29,37^ Taken together, these findings suggest an alternative pathway for DMT biosynthesis in rodents.

**Figure 1.**
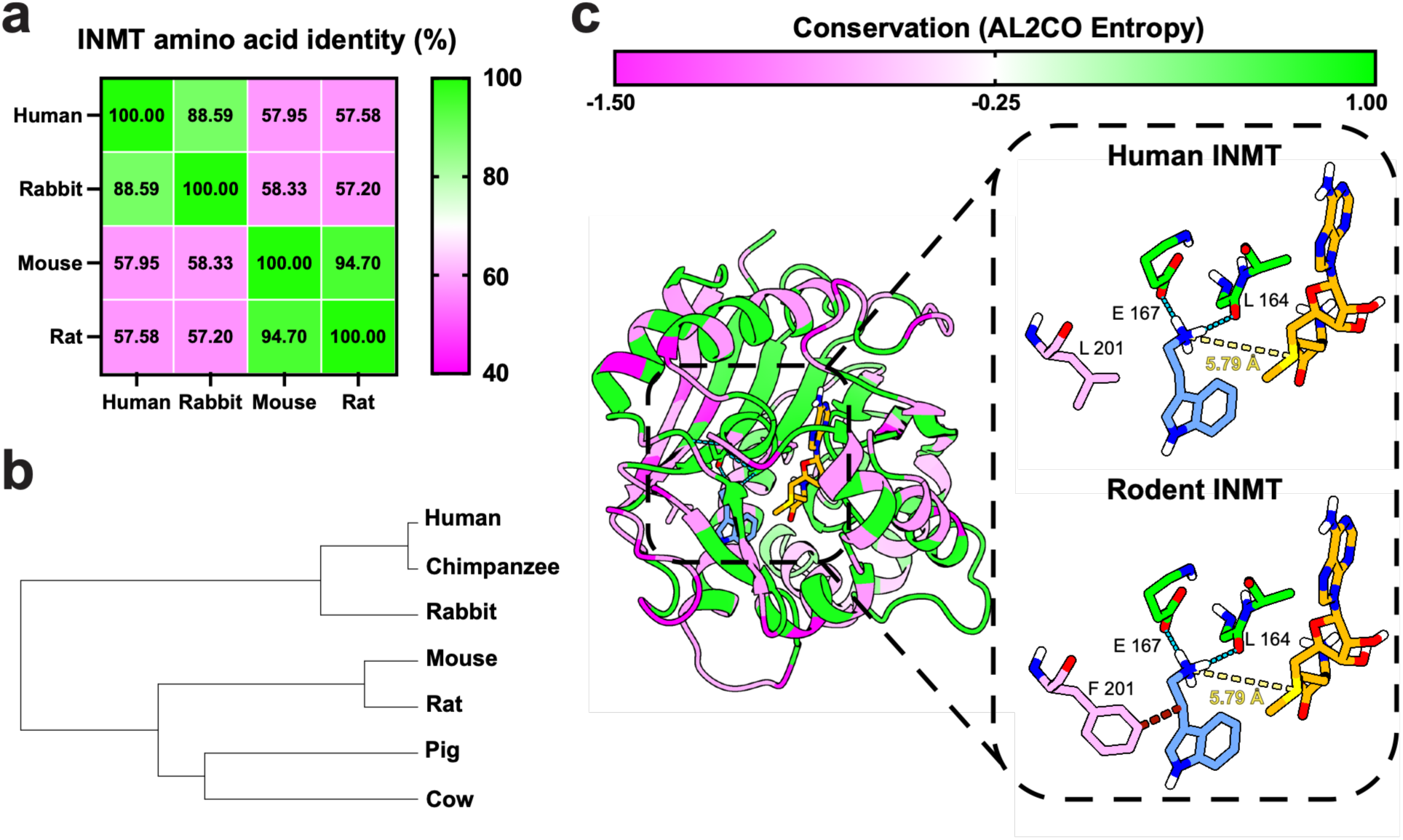
Comparative analysis of mammalian INMTs. **a)** Heatmap depicting the percent sequence identities between human, rabbit, mouse, and rat INMTs. Sequences were aligned using Clustal Omega and pairwise sequence identity was calculated in JalView. **b)** A neighbor-joining phylogenetic tree of human, chimpanzee, rabbit, mouse, rat, pig, and cow INMTs. JalView was used to generate the phylogenetic tree using the BLOSUM62 substitution matrix. **c)** Structural comparison of predicted tryptamine binding in human and rodent INMT catalytic sites. hINMT crystal structure (PDB: 2A14) was prepared with the co-crystallized *S*-adenosyl-L-homocysteine (SAH) cofactor modified to SAM and docked with protonated tryptamine. The L201F rodent variant was added to INMT *post hoc* to visualize effect on substrate binding. Residues are color-coded to illustrate calculated AL2CO entropy between human, chimpanzee, rabbit, mouse, rat, pig, and cow INMTs. Blue dashes represent hydrogen bonds; yellow dashes represent distances; red dashes represent steric clash.

Our docking analysis identified E167 as a putative catalytic base positioned to facilitate methyl transfer through the predicted hydrogen bond to the tryptamine substrate (**Figure 1c; Schematic S1**). Additionally, *in vitro* studies suggest that INMT activity is sensitive to redox state and product feedback inhibition, as DMT exhibits mixed inhibition relative to tryptamine.^38,39^ These enzymatic features support a model where INMT activity is regulated by both catalytic site geometry as well as cellular conditions, including redox state and product accumulation. Therefore, to assess the feasibility of local DMT biosynthesis in the brain, we need to elucidate INMT abundance and localization in human CNS cell types. We hypothesized that INMT localizes to defined subcellular compartments in human neurons to enable spatially-organized psychedelic tryptamine biosynthesis. To test this, we mapped INMT expression, subcellular localization, and topology in human induced pluripotent stem cell (iPSC)-derived neurons, a characterization not previously reported.

## RESULTS AND DISCUSSION

Until now, efforts to characterize endogenous psychedelic tryptamine biosynthesis in mammalian brain have relied on mRNA detection methods or *in vitro* enzymatic studies.^26,27,28,32,33,34,35,36^ To demonstrate INMT expression at the protein level in human neurons using an immunocytochemical (ICC) approach, we first validated a commercially available anti-INMT antibody. We evaluated the specificity of the antibody for the INMT epitope through an immunogen competition assay using human thyroid medullary cancer cell line TT cells (ATCC: CRL-1803), previously shown to express INMT.^40^ The anti-INMT antibody was pre-incubated overnight with 100-fold molar excess of its immunizing peptide before application to fixed TT cells along with a non-pre-incubated control. Fluorescence microscopy revealed a discrete and punctate INMT-derived signal in the non-pre-incubated condition that was abolished by peptide pre-incubation (**Figure S1**), confirming antibody reactivity with the INMT peptide. We extended this validation experiment to 6-week differentiated human iPSC-derived neurons, whereby the non-pre-incubated antibody showed a large population of punctate perinuclear INMT immunoreactivity, with smaller puncta above or on the nucleus and dendrites, all of which were ablated in the pre-incubated antibody condition (**Figure S2**). Taken together, these findings support the antibody’s ability to bind the INMT epitope and suggest that INMT spatially associates with subcellular compartments, rather than being distributed homogeneously in the cytoplasm.

To validate that the anti-INMT antibody recognizes the INMT epitope *in cellulo*, we generated a stable INMT-overexpressing HEK293T (ATCC: CRL-3216) cell line (HEK-INMT-OE) using an EF1α-driven lentiviral vector expressing hINMT (NCBI RefSeq ID: NM_006774.5) with a C-terminal FLAG® tag (VectorBuilder VB240605-1567zeq) (**Figure S3**). Following construct expression in wildtype HEK293T (WT HEK) cells and puromycin selection, fixed cells were co-incubated with the anti-INMT antibody and an anti-FLAG antibody and imaged using confocal fluorescence microscopy (**Figure 2a**). ImageJ (Fiji) was used to quantify the linear correlation between signals derived from the anti-INMT and anti-FLAG antibodies using the Bioimaging and Optics Platform within the Just Another Colocalization Platform (BIOP-JACoP). This analysis yielded a Pearson’s correlation coefficient (r = 0.94) between the two antibody-derived signals, supporting the conclusion that they likely detect the same protein (**Figure 2b**). Immunoblot analysis of HEK-INMT-OE cells using the anti-INMT and anti-FLAG antibodies both revealed a single ∼28 kilodalton (kDa) band at the predicted molecular weight (**Figure 2c,e**; full unedited membranes in Supporting Information, **Figure S4**). Densitometric quantification showed increased INMT and FLAG immunoreactivity in HEK-INMT-OE cells relative to the parental HEK293T cell line (**Figure 2d,f**).

**Figure 2:**
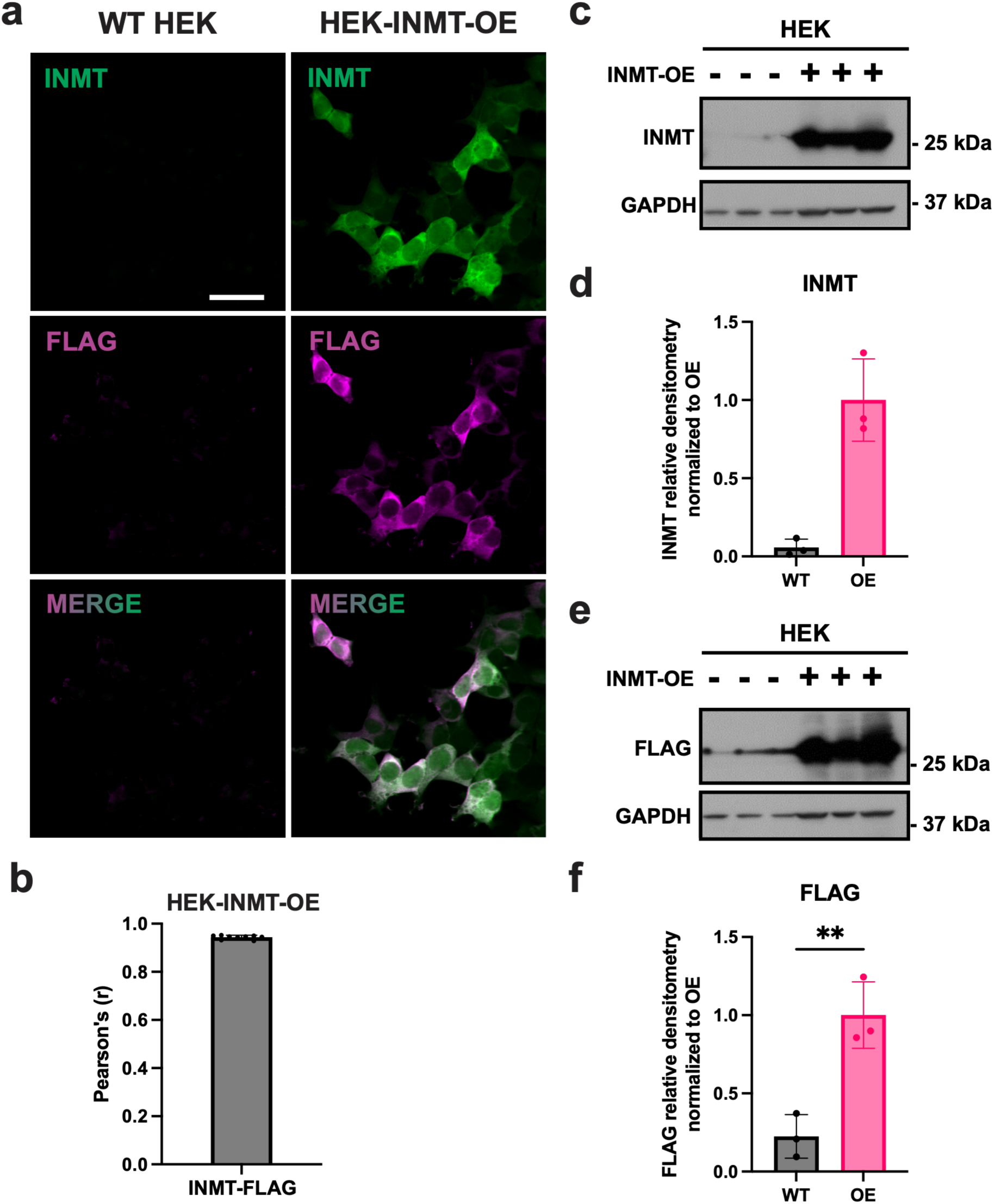
Overexpression of INMT in HEK293T cell line confirms antibody specificity. **a)** Representative images of HEK293T cell lines co-incubated with anti-INMT and anti-FLAG antibodies. *Left column:* WT HEK cells. *Right column:* HEK-INMT-OE cells. Scale bar: 10 μm. **b)** Quantification of spatial proximity as measured by Pearson’s r coefficient between INMT and FLAG immunoreactivity in HEK-INMT-OE cells. *n* = 9 images; 5 – 20 cells per image. One outlier was removed using ROUT in GraphPad Prism (*Q* = 1%). Bar represents mean ± SD, dots represent mean from a single image. **c)** Immunoblot illustrating INMT immunoreactivity in WT HEK and HEK-INMT-OE cells. **d)** Densitometric quantification of INMT immunoreactivity, relative to GAPDH, in WT HEK and HEK-INMT-OE cells. **e)** Immunoblot illustrating FLAG immunoreactivity in WT HEK and HEK-INMT-OE cells. **f)** Densitometric quantification of FLAG immunoreactivity, relative to GAPDH, in WT HEK and HEK-INMT-OE cells. Densitometric quantifications were normalized to the overexpression condition. Membranes developed using Pierce ECL Western Blotting Substrate. WT - WT HEK; OE - HEK-INMT-OE; GAPDH - Glyceraldehyde 3-phosphate dehydrogenase. Bars represent mean ± SD. *p < 0.1, **p < 0.01; unpaired Welch’s t test; *n* = 3 biological replicates.

As an orthogonal method of antibody validation, we performed RNA sequencing on the WT HEK and HEK-INMT-OE cell lines. Comparing transcripts between the two cell lines revealed only two significantly upregulated mRNAs, with *INMT* having over 10 log_2_ fold change in the HEK-INMT-OE cell line relative to the WT HEK cell line (**Figure S5**). The clear mRNA upregulation of *INMT* in the overexpression system suggests the increased INMT immunoreactivity observed in **Figure 2** can be attributed to increased INMT protein expression.

To further validate the specificity of the anti-INMT antibody, we performed a short hairpin RNA (shRNA)-mediated knockdown of INMT in HEK293T cells. HEK293T cells were transduced with lentiviral particles encoding a 29-mer shRNA targeting human *INMT* (NCBI Gene ID: 11185) or a scrambled control shRNA, both in a green fluorescent protein (GFP) reporter pGFP-C-shLenti vector (OriGene TL312141V). Colonies exhibiting a high degree of lentiviral integration were selected from experimental and control conditions to ensure comparable degrees of transduction (**Figure S6a**). Immunoblot analysis of both transduced cell lines revealed that the scramble shRNA control exhibited 2 bands at ∼28 kDa that were reduced in the INMT-targeting shRNA condition relative to the loading control (**Figure S6b-e**). This finding further suggests that the anti-INMT antibody recognizes INMT in a human cellular context.

*INMT* and *AADC* mRNAs were shown to exist in close proximity in rat brain, and DMT biosynthesis has been postulated to occur in distinct subcellular compartments to achieve physiologically relevant concentrations.^28,24,29^ We sought to expand this work by using protein-detection methods to detect INMT, AADC, and MAO-A to uncover if the spatial organization of DMT biosynthetic machinery is positioned to facilitate DMT production at specific subcellular compartments. Fixed human iPSC-derived neurons were co-incubated with the anti-INMT antibody in combination with previously validated anti-AADC or anti-MAO-A antibodies, as well as an antibody against microtubule-associated protein 2 (MAP2), a dendritic marker (**Figure 3a,b**).^41^ We used the BIOP-JACoP Fiji colocalization plugin in combination with our object-based segmentation pipeline (**Code S1**) to quantify the spatial proximity between INMT and AADC or MAO-A at single-cell resolution. In brief, the pipeline reports the linear correlation (Pearson’s r) and fraction of overlap (Manders’ M1 and M2) between two channels within a defined region of interest (ROI) that encapsulates a neuron’s soma and dendrites. The Manders’ M1 coefficient describes the fraction of pixels in channel A (INMT) above an intensity threshold (positive pixels) that are also positive in channel B (comparison protein), and the Manders’ M2 coefficient is the inverse. This analysis revealed that AADC demonstrated a significantly higher linear correlation with INMT than MAO-A (**Figure 3c**; median r = 0.43 and 0.30, respectively). Interestingly, we found no significant difference in M1 values (**Figure 3d**; INMT-AADC median M1 = 0.68; INMT-MAO-A median M1 = 0.63). AADC demonstrated a significantly greater overlap with INMT when compared to the overlap of MAO-A with INMT (**Figure 3e**; median M2 = 0.36, and 0.29, respectively). These data suggest that AADC, MAO-A, and INMT may occupy similar subcellular regions in human neurons, but AADC spatial intensity correlates more tightly with INMT. Proximity between INMT and AADC may support production of methylated tryptamines, while relative separation from MAO-A could limit product elimination by oxidative deamination. Taken together, the stronger proximal signals between AADC and INMT, combined with a significantly reduced INMT-MAO-A correlation, support the spatial feasibility of localized DMT biosynthesis.

**Figure 3:**
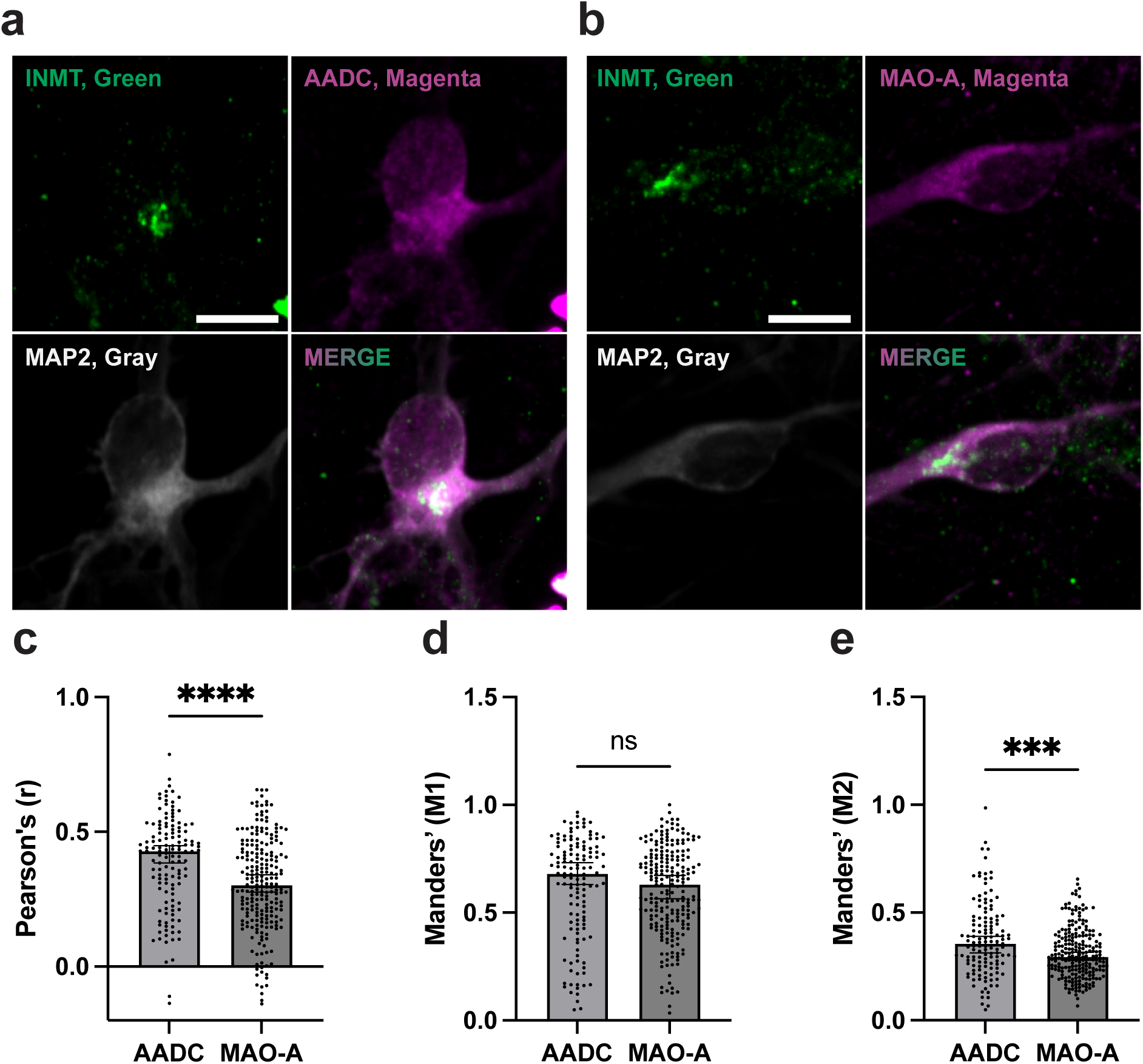
Single-cell spatial proximity analysis of INMT with AADC and MAO-A in human neurons. Representative images of human iPSC-derived neurons co-incubated with anti-INMT, anti-MAP2 and either **a)** anti-AADC or **b)** anti-MAO-A antibodies. Channel intensity was held constant within each antibody across images. **c)** Quantification of linear correlation (Pearson’s r) between INMT and AADC or MAO-A immunoreactivity. **d)** Quantification of fraction of overlap between INMT and AADC or MAO-A immunoreactivity. **e)** Quantification of fraction of overlap between AADC or MAO-A and INMT immunoreactivity. Bars represent median ± 95% CI; dots indicate individual neurons. AADC comparisons were derived from 133 individual cells; MAO-A comparisons were derived from 255 individual cells. INMT-negative cells were excluded from analysis. ns = not significant; ***p < 0.001; ****p < 0.0001; two-tailed Mann–Whitney U test. Scale bars: 10 μm.

After we determined that human neurons express a punctate, perinuclear subpopulation of INMT, we sought to identify the subcellular structure of INMT localization. Visually, this staining pattern resembled the Golgi apparatus, so we co-incubated fixed human iPSC-derived neurons with antibodies against INMT and Golgin subfamily A member 2 (GM130), a well-established *cis-*Golgi network marker (**Figure 4a**). To measure differences between INMT and GM130 peak intensities in two-dimensional space, we employed a line ROI-based protocol first described by van Bommel et al. (2023).^42^ In brief, this analysis is performed by selecting neurons where Golgi stacks are in line with the visual plane and drawing a linear ROI perpendicular to the Golgi stack starting from the *cis-*Golgi network and extending to the *trans*-Golgi network (see **methods**). To remove additional selection bias, only the GM130 marker was visualized when generating the ROI. We plotted INMT- and GM130-derived intensity along the ROI as a function of distance (**Figure 4b**) and calculated the median distance between INMT and GM130 maximum intensities averaged across all sampled cells (**Figure 4c**). These results revealed a median distance of 0 μm between INMT and GM130 intensity peaks, supporting the notion of INMT localization to the *cis*-Golgi network. As the *cis*-Golgi network receives trafficked cargo from the endoplasmic reticulum (ER) and functions as an early sorting and recycling station of the secretory pathway, localization of INMT to the *cis*-Golgi network may couple its catalytic activity to this key regulatory region of the secretory pathway where local pools of membrane-permeable tryptamines could form and influence signaling.

**Figure 4:**
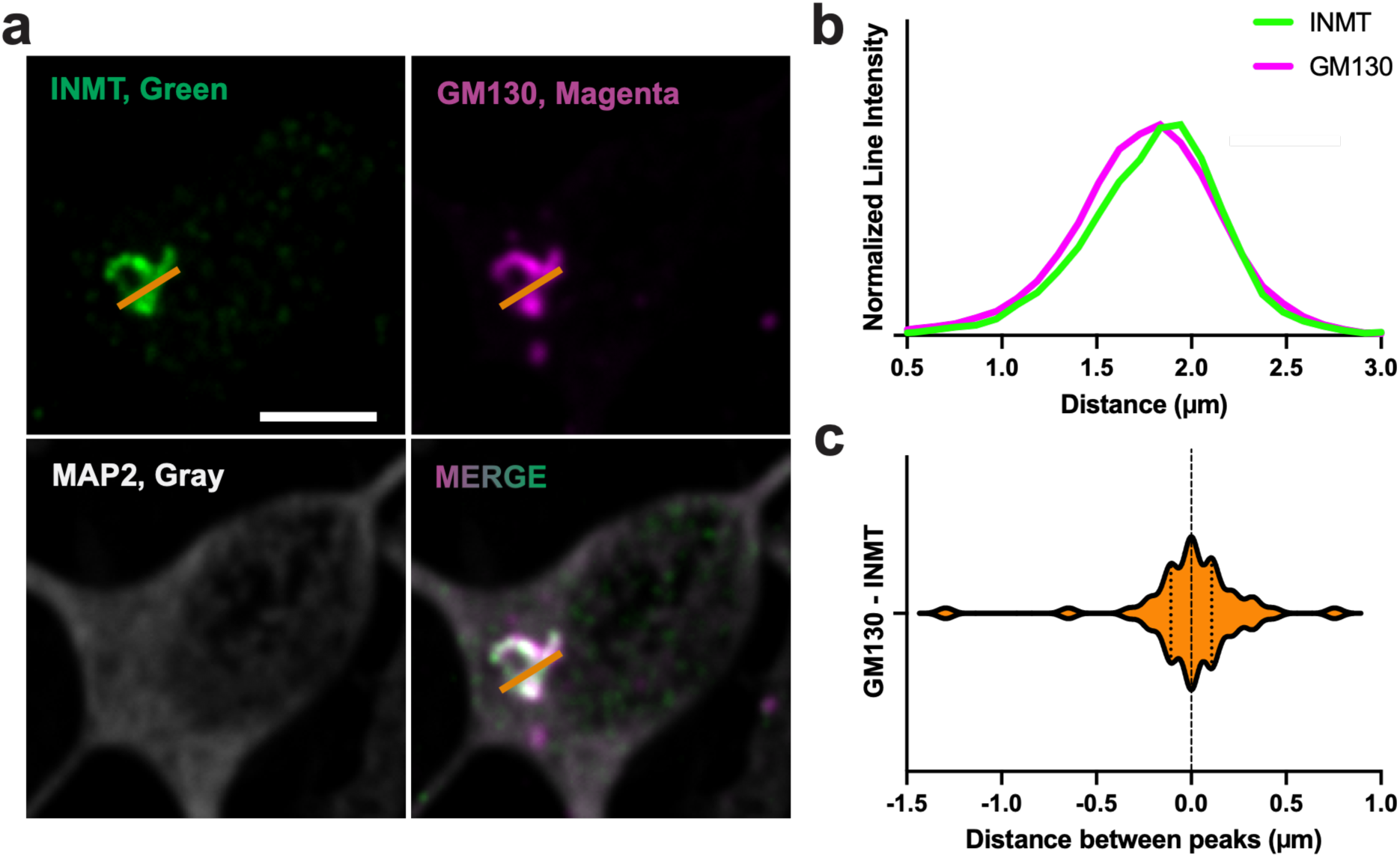
INMT localizes proximal to the *cis*-Golgi network. **a)** Representative images of human iPSC-derived neurons co-incubated with anti-INMT, anti-GM130, and anti-MAP2 antibodies. The orange line depicts an ROI used in downstream line intensity analysis. Scale bar: 5 µm. **b)** Representative intensity profile of INMT and GM130 immunoreactivity along the ROI shown in **a)**. Intensities were baseline-corrected and normalized before plotting. **c)** Distances between maximum intensity peaks of GM130 and INMT immunoreactivity. A violin plot represents the distribution of individual neurons. The dashed line represents the median; dotted lines show Q1 and Q3. *n* = 48 neurons.

While investigating INMT localization to the *cis-*Golgi network, we observed heterogeneous phenotypic profiles of Golgi structures. Recent work demonstrated that the Golgi apparatus is dynamic in human neurons and possesses various morphologies, and that this organelle trafficking is linked to neuronal polarization.^43^ To investigate Golgi phenotype subclasses in relation to INMT, fixed neurons were co-incubated with anti-INMT and anti-GM130 antibodies and imaged using confocal fluorescence microscopy. After imaging, each Golgi was assigned a phenotype using the 5 profile assignments previously described by Wang et al. (2023): compact somatic, soma-to-dendrite deployment, long filament, discrete ministack, and transient translocation (**Figure 5a**).^43^ We calculated the percentage of each phenotype across the neuronal population, revealing that 58% of neurons displayed a compact somatic phenotype (**Figure 5b**). The second most populous Golgi apparatus phenotype was soma-to-dendrite deployment at 15%, which represent a static snapshot of dendritic Golgi outpost generation. The next most populous subtype was transient translocation at 13%, a one-time event in neuronal differentiation wherein the entire Golgi apparatus leaves the soma and travels down a dendrite, after which neuronal polarity is fixed.^43^ Finally, the smallest Golgi subpopulations observed were dendritic phenotypes, with discrete ministack comprising 15% of the total population and long filament at just 5%. This population breakdown reflects what was previously described in human iPSC-derived neurons.^43^ To test whether INMT was enriched in a specific phenotype, we quantified average INMT intensity across Golgi phenotypes. The images were segmented using a GM130 mask and intensity of both INMT and GM130 immunoreactivities were recorded per GM130 object (see **methods**). To avoid biasing the selection towards more INMT-positive cells, Golgi were selected only using the MAP2 and GM130 channels for visualization. This analysis revealed no significant difference between any of the Golgi phenotypes (**Figure 5c**), establishing INMT as a putative Golgi apparatus resident protein.

**Figure 5:**
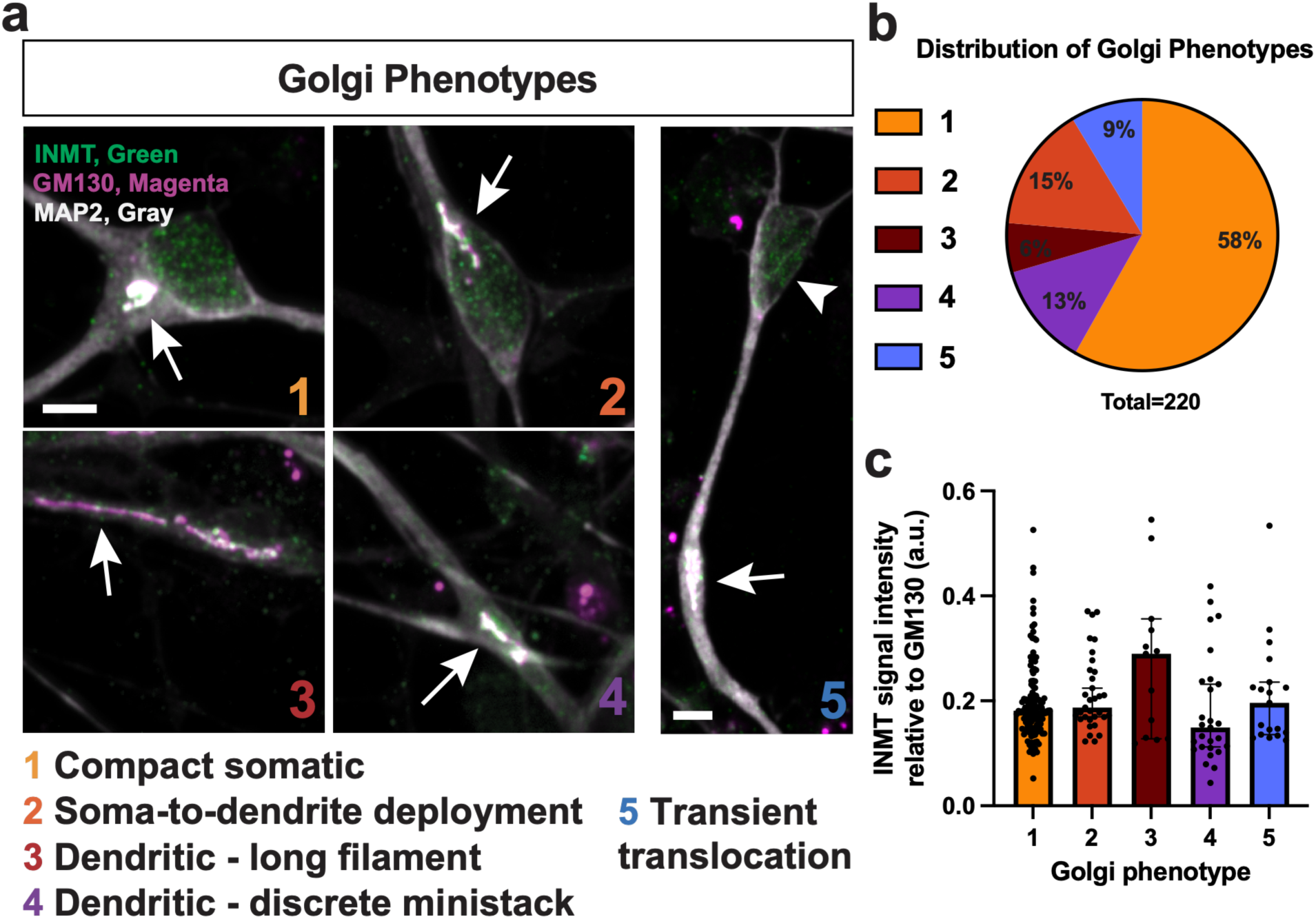
INMT presence across Golgi phenotypes in human iPSC-derived neurons. **a)** Representative images illustrating Golgi phenotypes present in human iPSC-derived neurons. Arrows highlight exemplar INMT-containing Golgi structures; an arrowhead highlights a soma without a Golgi apparatus. Channel intensity was held constant within each antibody across images. Scale bar: 10 µm. **b)** Percentages of each Golgi phenotype across a population of human iPSC-derived neurons. **c)** Single-cell intensity analysis of INMT immunoreactivity per Golgi phenotype. INMT immunoreactivity was normalized to GM130 immunoreactivity. Bars represent median ± 95% CI; dots represent mean immunoreactivity from a single ROI; *N* = 220 neurons; *n* = 13 - 128 neurons per group. No significant differences were detected across phenotypes by Kruskal–Wallis test followed by Dunn’s multiple-comparisons test.

In neurons, the Golgi apparatus overcomes spatial obstacles posed by complex neuronal morphology by trafficking Golgi structures from the soma to dendrites, where they serve as local secretory hubs nearer the synaptic cleft.^43^ Importantly, INMT immunoreactivity was not restricted to somatic Golgi structures but was also detected in discrete Golgi ministacks which are of particular interest as they are proposed to function as non-centrosomal microtubule-organizing centers.^44^ These myosin-dependent outposts support local protein trafficking and microtubule nucleation in distal dendrites, far from the parent somatic Golgi structures.^43,44^ Localization of INMT to these dendritic Golgi outposts suggests that INMT-facilitated tryptamine methylation may intersect with microtubule assembly and downstream structural plasticity. INMT was also detected at transiently translocated Golgi structures, a phenotype associated with human neuronal differentiation and characterized by deployment of the entire Golgi apparatus from the soma to a developing dendrite.^43^ This process occurs during neuronal polarization, when Golgi apparatus is linked to establishing cellular polarity.^43,45^ Demonstrating INMT’s presence near the Golgi apparatus during this polarization event suggests tryptamine methylation may be occurring at the Golgi during an event known to be implicated in neuronal maturation.

After establishing INMT localization in spatial proximity to the *cis*-Golgi network, we next examined whether INMT, DMT-like, and 5-HT_2A_ R immunoreactivity localize to the same subcellular regions in human neurons. Fixed neurons were co-incubated with previously validated antibodies against INMT, DMT-like small molecules, or the 5-HT_2A_ R together with anti-GM130, and anti-MAP2 antibodies, along with Hoechst nuclear stain for structural insight (**Figure 6a-c**).^46,20^ First, we quantified the spatial relationship between INMT, DMT-like, or 5-HT_2A_ R immunoreactivity and GM130 immunoreactivity using our single-cell object-based spatial proximity analysis pipeline (**Code S1**). Both INMT and DMT-like immunoreactivity exhibited a positive linear correlation with GM130, (INMT: r = 0.54; DMT: r = 0.33) (**Figure 6d**). In contrast, 5-HT_2A_ R immunoreactivity did not demonstrate a significant global linear correlation with the *cis*-Golgi network marker (r = 0.0) (**Figure 6d**). Despite low whole-cell associations between 5-HT_2A_ R and GM130 immunoreactivity, visually there appears to be a small population of 5-HT_2A_ R signal localized to GM130-positive structures (**Figure 6c**).

**Figure 6:**
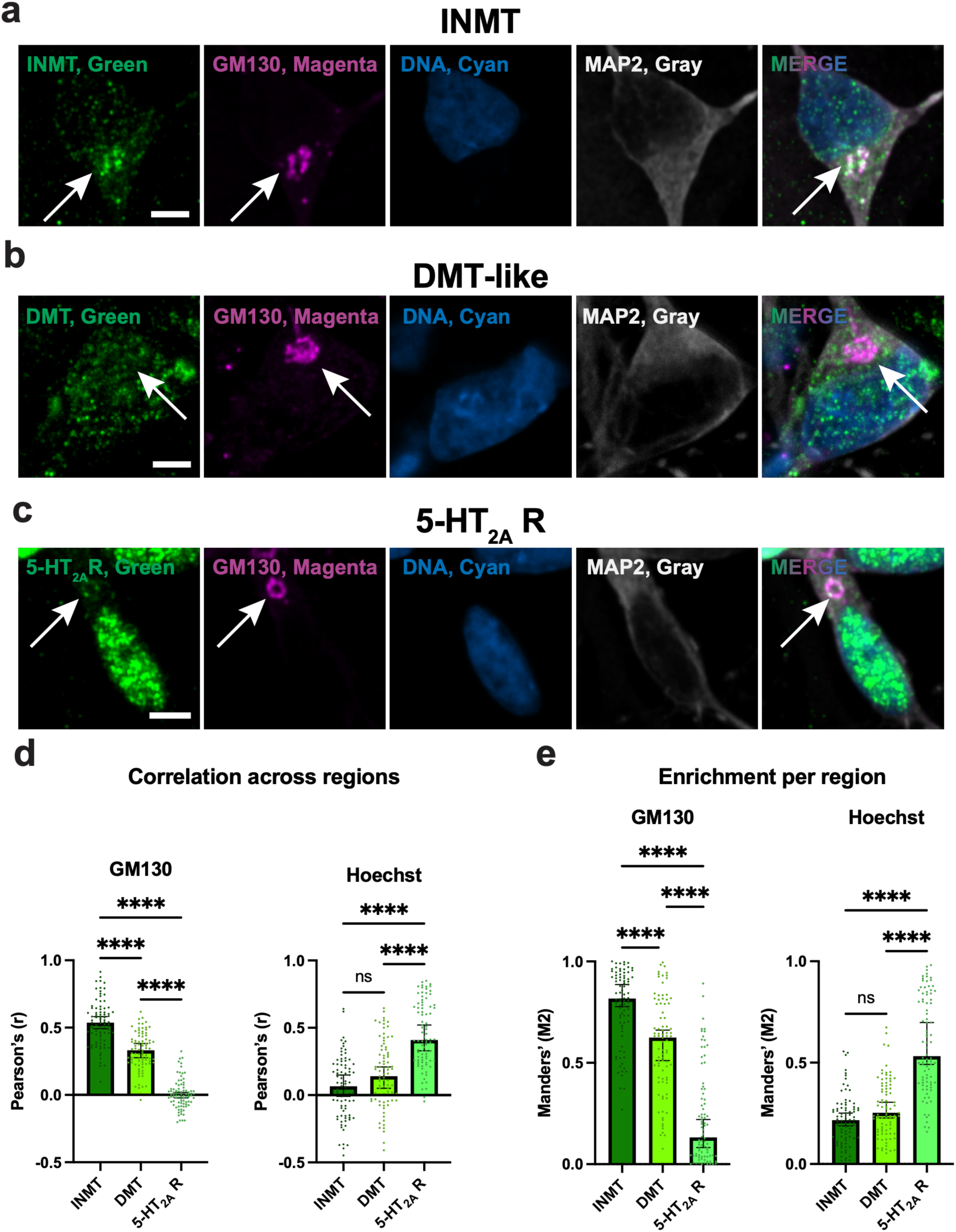
Single-cell Golgi and nucleus localization profiles of INMT, DMT-like, and 5-HT_2A_ R immunoreactivity. Representative images of human iPSC-derived neurons co-incubated with anti-GM130 and anti-MAP2 antibodies and Hoechst nuclear stain, together with either **a)** INMT, **b)** DMT-like, or **c)** 5-HT_2A_ R antibodies. White arrows highlight exemplar Golgi structures. Channel intensity was held constant within each antibody across images. **d)** Quantification of linear correlation (Pearson’s r) between DMT-related biomolecules and GM130 or Hoechst structural markers. **e)** Quantification of fraction of overlap (Manders’ M2) between GM130 or Hoechst structural markers and DMT-related biomolecules. Bars represent median ± 95% CI; dots indicate individual neurons. ns = not significant, ****p < 0.0001. Statistical comparisons among INMT, DMT, and 5-HT_2A_ R groups were performed separately for each marker using Kruskal–Wallis tests followed by Dunn’s multiple comparisons tests. *n* = 84 - 86 neurons per group. Scale bars: 5 μm.

5-HT_2A_ R immunoreactivity showed clear visual overlap with Hoechst-derived signal (**Figure 6c**), so we also quantified the linear correlation between each DMT-related biomolecule and the DNA marker (**Code S1**). Specifically, INMT and DMT-like immunoreactivity showed weakly positive association with Hoechst (INMT: r = 0.07; DMT: r = 0.14), while 5-HT_2A_ R immunoreactivity was strongly positively correlated (r = 0.41) (**Figure 6d**).

To capture the region-specific enrichment of each DMT-related biomolecule, rather than relying solely on whole-cell, intensity-sensitive Pearson analysis, we also quantified Manders’ M2 coefficients using our object-based spatial proximity analysis pipeline (**Code S1**). This metric represents how much of each subcellular compartment, defined by GM130-labeled Golgi structures or Hoechst-labeled nuclei, contains detectable INMT, DMT-like, or 5-HT_2A_ R immunoreactivity. INMT and DMT-like immunoreactivity showed high overlap with GM130 (INMT: M2 = 0.82; DMT: M2 = 0.62) (**Figure 6e**). 5-HT_2A_ R exhibited some overlap with GM130-positive pixels (M2 = 0.13), consistent with a Golgi-localized 5-HT_2A_ R subpopulation (**Figure 6e**). Examining the enrichment of INMT and DMT-like immunoreactivity at the nucleus revealed similarly low M2 values (INMT: M2 = 0.22; DMT: M2 = 0.25), while 5-HT_2A_ R showed greater overlap (5-HT_2A_ R: M2 = 0.53) (**Figure 6e**). Together, these findings support enrichment of INMT and DMT-like immunoreactivity at the Golgi apparatus and indicate that while 5-HT_2A_ R immunoreactivity most strongly localizes to the nuclear marker, there is a distinct subpopulation present near the *cis-*Golgi network, consistent with prior work in rodents describing a Golgi-associated 5-HT_2A_ R subpopulation.^47,48,49,50,20^

After establishing that INMT localizes to the *cis-*Golgi network in human neurons and 5-HT_2A_ R shows primarily nuclear localization, we sought to determine their topologies at their respective intracellular compartments. To distinguish cytosol-from lumen-facing epitopes, we performed an ICC-based selective permeabilization assay comparing immunoreactivity between Triton X-100 fully permeabilized conditions and digitonin partially permeabilized conditions. Under optimized conditions, digitonin selectively permeabilizes cholesterol-rich bilayers, like the plasma membrane, by forming water-insoluble complexes with cholesterol, leaving cholesterol-deficient membranes, like the Golgi membrane and nuclear envelope, intact.^50,52,53^ Under digitonin-permeabilized conditions, lumen-facing epitopes should not be antibody-accessible, while cytosol-facing epitopes should remain accessible.

To identify digitonin concentrations that permeabilize the plasma membrane but preserve Golgi membrane integrity, fixed human iPSC-derived neurons were treated with increasing concentrations of digitonin (6.25–50 μg/mL) along with a full permeabilization condition (0.3% Triton X-100). Next, neurons were co-incubated with antibodies against GM130, a cytosol-facing protein, and trans-Golgi network protein 46 (TGN46), a lumen-facing protein, along with an anti-MAP2 antibody (**Figure S7a**). TGN46 signal from each digitonin condition was measured per GM130-defined ROI in Fiji and normalized to signal intensity of the full permeabilization condition (**Figure S7b**; **Code S2**). We selected 25 μg/mL digitonin as the optimal concentration, as TGN46 immunoreactivity was diminished at this concentration relative to the full permeabilization condition.

Using this optimized digitonin condition, we next assessed INMT and 5-HT_2A_ R topology. Fully and partially permeabilized human iPSC-derived neurons were incubated with antibodies against either INMT or the 5-HT_2A_ R N-terminus, together with anti-MAP2 and anti-GM130 antibodies and Hoechst nuclear stain for structural compartmentalization (**Figure 7a&b**). Images were analyzed using an ROI-based Fiji code that reports on mean signal intensity of either INMT or 5-HT_2A_ R within GM130- or Hoechst-defined subcellular regions (**Code S2**).

**Figure 7:**
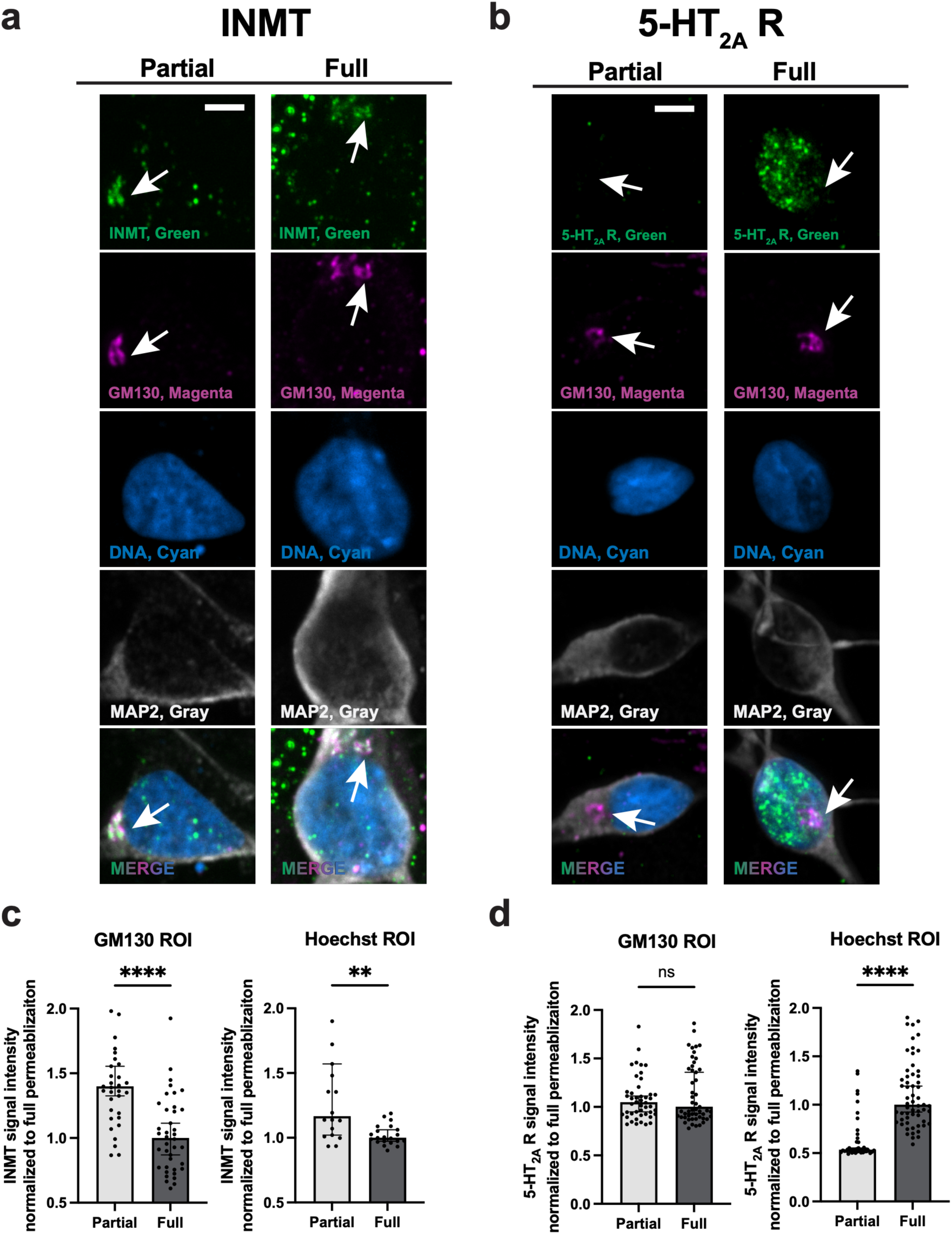
Selective permeabilization analysis of INMT and 5-HT_2A_ R epitope accessibility. Representative images of human iPSC-derived neurons co-incubated with antibodies against either **a)** INMT or **b)** 5-HT_2A_ R N-terminus, along with anti-GM130 and anti-MAP2 antibodies and Hoechst nuclear stain. White arrows highlight exemplary Golgi structures. The left and right columns show results from partial and full permeabilization conditions, respectively. Channel intensity was held constant within each antibody across images, except for GM130, which varied between permeabilization conditions for visual clarity. Quantification of **c)** INMT and **d)** 5-HT_2A_ R immunoreactivity intensity from both partial and full permeabilization experimental conditions, normalized to the full permeabilization condition. For both **c)** and **d)**, left graphs depict the normalized intensity measured within the GM130 ROI, and right graphs depict the normalized intensity measured within the Hoechst ROI. Bars represent median ± 95% CI; dots indicate individual ROIs. *n* = 18 – 59 ROIs per group. ns = not significant; **p < 0.01, ****p < 0.0001; two-tailed Mann-Whitney U test. Scale bars: 5 μm.

INMT immunoreactivity at the GM130-defined region under partial permeabilization was increased relative to the full permeabilization condition, which is consistent with a cytosol-facing topology at the Golgi membrane (**Figure 7c**). This increase may represent improved native epitope preservation relative to the harsher Triton X-100 permeabilization condition. When we assessed changes in INMT immunoreactivity in the Hoechst-defined compartment, INMT also showed a significant increase under partial permeabilization, consistent with a cytosol-facing topology at the nucleus (**Figure 7c**).

In contrast, 5-HT_2A_ R immunoreactivity within the GM130-defined region showed no statistical difference between the two permeabilization conditions but was significantly reduced within the Hoechst-defined region in the partial permeabilization condition, consistent with an intraorganelle-facing N-terminus (**Figure 7d**). The absence of a significant reduction in immunoreactivity within GM130-defined regions may reflect the relatively small and heterogeneous Golgi-associated 5-HT_2A_ R population identified above, as many GM130-positive regions contained little or no detectable 5-HT_2A_ R immunoreactivity. Thus, any permeability-dependent loss of Golgi-associated 5-HT_2A_ R signal may be diluted when quantified across all GM130-positive regions. The significant reduction in Hoechst-associated 5-HT_2A_ R immunoreactivity under partial permeabilization is consistent with restricted antibody access of the N-terminal receptor epitope, supporting an intraorganelle-facing binding pocket for this receptor population, consistent with biological convention.

The 5-HT_2A_ R desensitizes in response to agonist binding and subsequently internalizes to early endosomes, where it was historically thought to be recycled or degraded.^54,55,56^ Alternatively, modern interpretations propose that internalized GPCRs can engage signaling pathways at intracellular compartments that are distinct from those activated at the plasma membrane, suggesting that 5-HT_2A_ R internalization could fulfil a biological role.^17,18,19^ Indeed, previous work demonstrated the ability of DMT and bufotenine to passively diffuse across cellular membranes and activate an intracellular 5-HT_2A_ R population, which resulted in increased structural plasticity not observed upon administration of hydrophilic 5-HT_2A_ R agonists unless aided by electroporation.^20^ This finding has led to an emerging hypothesis that DMT or bufotenine may be endogenous ligands of internalized 5-HT_2A_ Rs due to a superior passive ability to access this receptor pool over serotonin.^20,57^ Our localization of INMT, and subpopulations of 5-HT_2A_ R and DMT-like immunoreactivity to the Golgi apparatus supports this native ligand hypothesis by placing DMT biosynthetic machinery and potentially DMT itself within the same subcellular space. Future work should interrogate if local psychedelic tryptamine biosynthesis at the *cis*-Golgi network contributes to an agonism feedback loop, trapping the internalized 5-HT_2A_ R population where its intracellular signaling could influence cargo trafficking from the ER to the Golgi apparatus, and secretory pathway output.

Our discovery that INMT possesses cytosol-facing topology at the Golgi apparatus has important implications for both substrate access and enzymatic regulation. Positioning INMT on the cytosolic face of the *cis*-Golgi network would enable the enzyme to access AADC-synthesized tryptamine present in the cytosol without requiring active transport across the Golgi membrane. INMT’s cytosol-facing topology is likely functionally important, as the cytosol is more reducing than the Golgi lumen and INMT shows increased catalytic efficiency under reducing conditions *in vitro*.^58,59,39^ Moreover, cellular processes and disease states associated with reductive stress may augment INMT activity and affect downstream signaling pathways *in vivo*. Acute hypoxia generates reductive stress and, under this mechanistic framework, could contribute to increased INMT-mediated production of methylated tryptamines.^60^ This hypothesis is supported by the observation that carbon dioxide-induced cardiac arrest in rats leads to an increase in the accumulation of endogenous tryptamines.^28^ Additionally, in a Sigma-1 receptor-dependent manner, administration of exogenous DMT protects against hypoxia-induced cell death in human neurons, reduces ischemic lesion size, and improves functional recovery in the rat cortex.^61,62^ Taken together, these findings suggest that hypoxia may upregulate psychedelic tryptamine biosynthesis, and potentially serve as an endogenous mechanism for neuroprotection against cardiac arrest and acute brain injury.

Our findings are consistent with biological convention that positions the Golgi-localized 5-HT_2A_ R binding pocket facing the lumen, indicating that small-molecule agonists must cross the Golgi membrane to access this receptor pool.^63^ The population of unprotonated methylated tryptamines that passively diffuse across the Golgi membrane is expected to become predominantly protonated once inside the lumen, as their basic amine groups readily accept protons under acidic luminal conditions (pH ∼6.4).^64^ Once protonated, these species are less membrane permeable and may accumulate within the acidic cellular compartment.^65^ This mechanism has been proposed to explain sustained therapeutic benefits of exogenous administration of psychedelic tryptamines after a single dose, long after compound clearance.^20,64,66,11,10^ A lumen-facing 5-HT_2A_ R binding pocket, coupled with location bias theory and acid trapping could extend agonist availability and lead to prolonged activation of this receptor pool. This phenomenon also addresses a central concern often raised when establishing DMT biosynthesis in the human CNS: whether DMT can reach physiologically active concentrations.^67^ Taken together, our results suggest a model where INMT catalytic activity at the cytosolic face of the *cis*-Golgi network is positioned to create a local microdomain of membrane-permeable psychedelic tryptamines that can access an intracellular 5-HT_2A_ R pool.

INMT localization in close proximity to the cytosolic face of the *cis*-Golgi network opens the possibility that disruption of Golgi architecture could alter endogenous psychedelic tryptamine biosynthesis. Golgi dysfunction is implicated in a host of neurodegenerative disorders, and Golgi fragmentation is a well-established pathological feature of Alzheimer’s disease.^68,69^ Golgi fragmentation could cause INMT mislocalization, disrupt its fine-tuned catalytic activity, and contribute to downstream disease phenotypes.^70^ Taken together, INMT presence across neuronal Golgi subtypes suggests that INMT may participate in Golgi-mediated processes involved in regulating structural plasticity and neuronal development, and Golgi fragmentation could alter INMT function in neurodegenerative disease states. The role of endogenous psychedelic tryptamine biosynthesis and signaling may represent an underexplored avenue in human neurobiology, potentially with broad relevance to neurodevelopment, neuroplasticity, and disease.

## METHODS

### Species alignment

The NCBI database was used to acquire INMT amino acid sequences for: human isoform 1 (NP_006765.4), chimpanzee (XP_527713.4), rabbit (NP_001075512.1), mouse (NP_033375.1), rat (NP_001102492.1), pig (NP_001231662.1), and cow (NP_001179577.1). These sequences were aligned using Clustal Omega through JalView. After alignment, JalView was used to calculate the pairwise sequence identity between species, and a neighbor-joining phylogenetic tree was generated using a BLOSUM62 substitution matrix. The resulting tree was uploaded to Interactive Tree of Life for final edits. The aligned sequences were uploaded to ChimeraX and per-residue conservation scores were calculated using AL2CO entropy metrics. The resulting entropy scored were mapped onto hINMT (PDB: 2A14) and color-coded from magenta (high entropy) to green (low entropy) for visualization in **Figure 1**.

### Molecular Operating Environment *in silico* docking studies

Structural modeling and docking analyses were computed in Molecular Operating Environment (MOE, Chemical Computing Group). Human INMT (PDB: 2A14) was imported into the software and prepared using Protonate 3D at pH 7.2. The co-crystalized SAH cofactor was converted to SAM using the builder tool by addition of a methyl group to the sulfonium. Residues within 4.5 Å of SAM were subjected to local energy minimization to accommodate the modified cofactor. Putative substrate bindings sites near SAM were identified using the Site Finder module. The binding site included the following residues: Y20, Y24, Y25, L164, E167, C168, T198, L201, Y204, C213, Y242, A247. Tryptamine in its protonated state was docked using the Triangle Matcher placement method and refined using a rigid receptor model at the identified substrate binding pocket. Initial placement was scored using London dG and poses were refined using GBVI/WSA dG scoring. SAM was included in the receptor for docking. Pose shown in **Figure 1** was selected based on a catalytically competent orientation, defined by proximity of the substrate amine lone pair to the SAM methyl donor consistent with a concerted nucleophilic substitution reaction mechanism (S_N_2). A rodent INMT model was generated by introducing the L201F substitution, which is present in both rat and mouse INMTs, *post hoc* into the hINMT crystal structure. All other residues at the identified binding pocket are conserved between human and rodent INMTs. Structural comparisons between human and rodent models were used to assess the impact of this substitution on substrate accessibility of the binding pocket.

### TT cell culture

TT cells (ATCC CRL-1803) were cultured in T75 flasks (Corning 353136). TT cells were supplied with TT media consisting of 90% Ham’s F-12K Medium (Gibco 21127), 10% heat-inactivated fetal bovine serum (FBS) (GeminiBio 100-106), 1x penicillin/streptomycin (Gibco 15140) and filtered through a 0.22 μm filter (Nalgene 127-0020) prior to use. Cells received a 100% media change two times a week. Once cells reached confluency, cells were passaged using 1.5 mL TrypLE Select (Gibco 12563-029) per flask until detached. 10 mL of TT media was added to each flask and cell suspensions were centrifuged in a 15 mL conical tube (Falcon 14-959-49B) at 300 rcf for 5 minutes. The supernatants were aspirated and resuspended in TT media and seeded at a 1:3 dilution for expansion, or as described in following sections for downstream applications.

### HEK293T cell culture

HEK293T cells were cultured in T75 flasks (Corning 353136). HEK293T cells were supplied with basic growth media (BGM) consisting of 85% Dulbecco’s Modified Eagle Medium (DMEM) (Gibco 11995), 10% heat-inactivated fetal bovine serum (FBS) (GeminiBio 100-106), 1x penicillin/streptomycin (Gibco 15140) and filtered through a 0.22 μm filter (Nalgene 127-0020) prior to use. Cells received a 100% media change two times a week. Once cells reached confluency, cells were passaged using 1.5 mL TrypLE Select (Gibco 12563-029) per flask until detached. 10 mL of BGM was added to each flask and the cell suspensions were centrifuged in a 15 mL conical tube (Falcon 14-959-49B) at 300 rcf for 5 minutes. The supernatants were aspirated and resuspended in BGM and seeded at a 1:5 dilution for expansion, or as described in following sections for downstream applications.

### iPSC-derived neural progenitor cell culture

Human neural progenitor cells (NPCs) were generated from reprogrammed iPSCs derived from an individual with no known neuropsychiatric illness, as described by Sheridan et al., 2011 (GM08330) (8330–8, Coriell Institute for Medical Research, Camden, NJ).^71^ The generation of NPCs from these iPSCs was performed as described by Cheng et al., 2017.^72^ In brief, NPCs were cultured in 6-well plates (Falcon 353046 or CellTreat 229105) which were first coated with POL coating which consisted of 20 µg/mL poly-ornithine (Sigma Aldrich P35307) in ddH_2_O for 2 - 6 hours at 37 °C and 5% CO_2_, and next with 5 μg/mL laminin (Sigma Aldrich L2020) in PBS (Corning 21-040) overnight at 37 °C and 5% CO_2_. Plates were stored at 4 °C and used within 4 weeks.

NPCs were cultured in NS media consisting of 70% Dulbecco’s Modified Eagle’s Medium (DMEM) (Gibco 11995), 30% Ham’s F12 with L-glutamine (Modified Cellgro/Mediatech 10–080-CV), with 1x penicillin/streptomycin (Gibco 15,140–122), and 1x B27 Supplement (Gibco 17,504–044). On the day of use, the media was supplemented with 20 ng/mL Epidermal Growth Factor (EGF) (Sigma Aldrich E9644), 20 ng/mL basic Fibroblast Growth Factor (bFGF) (ReproCELL 03-0002), and 5 μg/mL heparin (Sigma Aldrich H3149), generating NS++ media. Once cells reached confluency, cells were passaged using 0.5 mL TrypLE Select (Gibco 12563-029) per well until detached. 1 mL of NS++ was added to each well, and the cell suspensions were centrifuged in a 15 mL conical tube (Falcon 14-959-49B) at 300 rcf for 5 minutes. The supernatants were aspirated and resuspended in NS++ media and seeded at a 1:3 dilution for expansion, or as described in the following sections for downstream applications.

### Human iPSC-derived neuron culture and differentiation

NPCs were plated in either a 96-well plate (Ibidi 89621) or 8-well chamber slide (Millipore PEZGS0816) precoated with POLS, consisting of 20 µg/mL poly-ornithine (Sigma Aldrich P35307) and 5 μg/mL laminin (Sigma Aldrich L2020) in PBS (Corning 21-040), overnight at 37 °C and 5% CO_2_. Upon the day of seeding, the plate or slide was aspirated to remove the POLS, and wells received 200 μL of NS media without growth factors per well. NPCs were passaged using 0.5 mL TrypLE Select (Gibco 12563-029) per well until detached. 1 mL of NS media without growth factors was added to each well, and the cell suspensions were centrifuged in a 15 mL conical tube (Falcon 14-959-49B) at 300 rcf for 5 minutes. The supernatants were aspirated and resuspended in NS media and seeded at 20,000 – 25,000 cells per well in 200 μL per well for a total volume of 400 μL per well. Cells were allowed to differentiate for 6 weeks at 37 °C and 5% CO_2_ and received half-media changes twice a week.

### Generation of HEK-INMT-OE cell line

HEK293T cells were cultured as described above. The day before transduction, cells were seeded in 6-well plates at a density of 300,000 cells per well and incubated at 37 °C with 5% CO_2_. hINMT (NM_006774.5) was cloned into a lentiviral expression vector under the EF1α promoter with a C-terminal FLAG tag (pLV-EF1α-hINMT-FLAG-Puro; VectorBuilder, VB240605-1567zeq). Viral stocks were diluted in BGM to achieve a working concentration of 3 x 10^6^ infectious units (IU) per mL. 1 mL of the viral media was added to each well to achieve a multiplicity of infection of 10. The day after viral transduction, wells received a 100% media change with BGM. 48 hours after transduction, puromycin (Gibco A1113803) was added to cellular media at a working concentration of 2 μg/mL, and cells were cultured under puromycin selection for 6 days. After puromycin selection, the HEK-INMT-OE cell line was cryopreserved, passaged, or pelleted for downstream applications.

### shRNA-mediated knockdown of INMT in HEK293T cells

HEK293T cells were cultured as described above. The day before transduction, cells were seeded in a 24-well plate (Falcon 353047) at a density of 100,000 cells per well and incubated at 37 °C with 5% CO_2_. The next day, cells were transduced using lentiviral particles encoding a pooled mixture of INMT-targeting shRNA constructs (OriGene, TL312141; Gene ID: 11185), each consisting of a unique 29-mer sequence cloned into a pGFP-C-shLenti backbone. A non-targeting scrambled shRNA control (OriGene, TR30021) in the same pGFP-C-shLenti backbone was used as a control. Viral stocks were added directly to cell media to achieve a multiplicity of infection of 15 for both the INMT-targeting and scramble shRNAs. The day after viral transduction, wells received a 100% media change with BGM. GFP expression began 48 hours after transduction. 12 days post-transduction, cells were passaged into a 96-well plate at a limiting dilution for clonal isolation. The highest GFP-expressing colonies were selected for downstream applications.

### General immunocytochemistry protocol

Unless stated otherwise in the following sections, this is the universal protocol for all ICC experiments. Cells were fixed by adding an equal volume of 8% paraformaldehyde solution (Tousimis 1008A) in PBS (Corning 21-040) directly to culture media to achieve a final concentration of 4% paraformaldehyde, and incubated for 1 hour at room temperature. The wells were washed 4x with 200 μL PBS (Corning 21-040) and subsequently incubated with 200 μL of blocking buffer consisting of 0.3% Triton-X 100 (Promega H5141), PBS (Corning 21-040) and 2% bovine serum albumin (Sigma Aldrich A7906) for 1 hour at room temperature. The blocking buffer was removed and cells were incubated with primary antibodies overnight at 4 °C, diluted in blocking buffer at the dilutions listed in **Table S1**, unless otherwise noted. After incubation with the primary antibody solution, wells were washed 4x with 200 μL PBS (Corning 21-040) and incubated with a 1:5000 dilution of Hoechst DNA stain (Invitrogen H3570) and secondary antibodies in blocking buffer at the dilutions listed in **Table S2** unless otherwise noted, for 1 hour at room temperature. Next, wells were washed 4x with 200 μL PBS (Corning 21-040) and the plate was sealed using aluminum sealing tape (Corning 6570). Fluorescence microscopy images were obtained using an InCell 6000 (GE Healthcare) automated microscope, a CellVoyager CQ1 (Yokogawa) spinning disk confocal microscope, or an AxioScan (Zeiss) digital slide scanner. Laser power and exposure times were identical within a set of experiments. Images intended for direct comparison were acquired using identical acquisition settings.

### Immunogen competition assay

TT cells were plated in an 8-well chamber slide (Millipore PEZGS0416) at a density of 40,000 cells/well. iPSC-derived human neurons were plated in a 96-well plate (Ibidi 89621) at a density of 12,000 cells/well. Both cell types were cultured according to protocols outlined above. Cells were fixed and permeabilized according to the general ICC protocol. The anti-INMT antibody was incubated overnight at 4 °C with 100x molar excess of its immunizing peptide. 20 μL of anti-INMT antibody (Novus NBP2-57073) (100 μg/mL) was added to a 1.5 mL conical tube. Next, 18.18 μL of the antigen peptide (Novus NBP2-57073PEP) (1,700 μg/mL, 1 M urea in PBS) was added to the same microcentrifuge tube. The total volume was brought to 40 μL with 1.9 μL PBS (Corning 21-040). This was repeated for the non-pre-incubated control with 18.18 μL of a 1 M urea (Sigma Aldrich U4883) solution in PBS (Corning 21-040) substituting the immunizing peptide. Both pre-incubated and non-pre-incubated controls were incubated overnight on an orbital shaker at 4 °C. The next day, 100 μL of PBS (Corning 21-040) was added to each antibody condition and wells were aspirated completely and incubated in blocking buffer according to the general protocol. Each well received 50 μL of either primary antibody mixture and was incubated at room temperature for 3 hours. Following primary antibody incubation, washes and secondary antibody incubation was performed according to the general ICC protocol using an anti-rabbit Alexa Fluor 488 antibody at the dilution listed in **Table 2**. Chamber slides were mounted using ProLong Mountant with DAPI (Invitrogen P36931) and a #1.5 coverslip (Fisher Scientific 12-541-016). Chamber slides were imaged using an AxioScan (Zeiss) digital slide scanner. All laser settings were consistent between conditions and resulting images were from a single focal plane. 96-well plates were imaged using an InCell 6000 automated microscope and images were from a single focal plane. Representative images all are shown at the same brightness and contrast levels set in Fiji.

### Immunocytochemistry of HEK-INMT-OE cell line

HEK-INMT-OE and WT HEK cells were passaged according to the HEK cell culture protocol and seeded into a 96-well plate (Corning 3904) at a density of 40,000 cells per well. Fixation and washing steps were performed as described in the general ICC protocol. Primary antibody incubation was performed overnight at 4 °C as described in the general ICC protocol, using anti-INMT and anti-FLAG antibodies at the dilutions listed in **Table S1**. Following primary antibody incubation, washes and secondary antibody incubation were performed according to the general ICC protocol using anti-rabbit Alexa Fluor 488, anti-mouse Alexa Fluor 647 antibodies, and Hoechst (Invitrogen H3570) at the dilutions listed in **Table 2**. 96-well plates were imaged using an InCell 6000 automated microscope and images analyzed downstream were from a single focal plane. Laser power and exposure times were identical between images within a set of experiments. Representative images all are shown at the same brightness and contrast levels set in Fiji.

### Immunoblotting and analysis

Cell pellets were collected via cell scraping of individual wells of a 6-well plate (CellTreat 229105). Cell suspensions were centrifuged at 1500 rpm in a 1.5 mL microcentrifuge tube, and supernatants were aspirated. Cell pellets were stored at -80 °C until use. 10 μL of each respective cell pellet was lysed directly in 80 μL of 1.5x SDS loading buffer (New England BioLabs B7703S) supplemented with 1.5x DTT (New England BioLabs B7705S) via vigorous reconstitution and incubation at room temperature for 15 minutes. Viscous samples were sonicated for 10 minutes. Lysates were then boiled at 95 °C for 10 min, briefly centrifuged, and allowed to equilibrate for 5 min. Proteins were separated on Tris-Glycine 4-20% gels (Invitrogen WXP42020BOX) in 1x Tris-Glycine SDS running buffer (Boston BioProducts BP-150). Each well received 10 μL of sample and gels were run at 125V for 2 hours. Gels were then transferred onto 0.45 μm polyvinylidene fluoride (PVDF) membranes (ThermoFisher PI88518) in transfer buffer (1x transfer buffer (Boston BioProducts BP-190), 20% methanol (Sigma-Aldrich 179337) in ddH_2_O) at 100 V for 1.5 hours. Membranes were blocked in blocking buffer consisting of 5% w/v BSA (Sigma-Aldrich A4737) in TBST (Boston BioProducts IBB-181X) for 1 hour at room temperature. Membranes were probed overnight at 4 °C with primary antibodies in blocking buffer at the dilutions listed in **Table S1**, unless otherwise noted. The following day, membranes were washed 3x for 10 minutes each with TBST (Boston BioProducts IBB-181X) and incubated for 1 hour at room temperature with species appropriate HRP-conjugated secondary antibodies in blocking buffer at the following dilutions listed in **Table S2** unless otherwise noted. Then, membranes were washed 3x for 10 minutes each with TBST (Boston BioProducts IBB-181X) and developed with chemiluminescence reagents (Pierce ECL Western Blotting Substrate, ThermoFisher 32106; SuperSignal West Pico Extended Duration Substrate, ThermoFisher 34577) on autoradiographic films (Labscientific XAR ALF 1318). Films were scanned on an Epson Perfection V600 Photo Color Scanner (Epson B11B198011). Membranes were stripped using 15 mL of stripping buffer (Life Technologies 46430) at room temperature for 15 minutes. Membranes were re-blocked in blocking buffer for 1 hour at room temperature reprobed for the loading control using an anti-GAPDH primary antibody (**Table S1**) under the same conditions as described above. After primary antibody incubation, membranes were incubated with the appropriate HRP-conjugated secondary antibody (**Table S2**), washed, developed, and imaged as described above. Bands were quantified using Fiji and normalized to GAPDH.

### Immunocytochemistry and confocal fluorescence microscopy for spatial proximity analysis and Golgi apparatus phenotyping

Human iPSC-derived neurons were differentiated according to the differentiation protocol described above. Fixation, permeabilization, blocking, and washing steps were performed as described in the general ICC protocol. Primary antibody incubation was performed overnight at 4 °C as described in the general ICC protocol using anti-INMT, anti-AADC, anti-MAO-A, anti-MAP2, anti-GM130, anti-TGN46, anti-5-HT_2A_ R, and anti-DMT-like antibodies at the dilutions listed in **Table S1**. The anti-5-HT_2A_ R antibody was previously validated in an immunocytochemical approach using INMT knockout mouse cortical neurons.^20^ The anti-DMT-like antibody was generated from rabbit antisera immunized with a BSA-conjugated DMT-derived haptogen and previously characterized by ELISA, demonstrating a limit of detection of 0.017 ± 0.005 ng/mL and <4% cross-reactivity across 101 psychoactive compounds, with psilocin showing the highest cross-reactivity at 3.6%.^46^ Following primary antibody incubation, secondary antibody incubation was performed according to the general ICC protocol using anti-rabbit Alexa Fluor 488, anti-mouse Alexa Fluor 647, anti-chicken Alexa Fluor 594, anti-sheep Alexa Fluor 488, and Hoechst at the dilutions listed in **Table S2**. Confocal fluorescence microscopy images were generates using a CellVoyager CQ1 (Yokogawa) spinning disk microscope. All images shown are maximum projections generated from Z-stack acquisition. Laser power and exposure times were held constant within a set of experiments. Representative images may have different brightness and contrast settings across channels to highlight subcellular structures.

### Digitonin-based permeabilization experiments

Human iPSC-derived neurons were differentiated for 6 weeks through the growth factor withdrawal protocol described above. The cells were fixed according to the general ICC protocol.

For the optimization experiment: Digitonin was serial diluted in PBS (Corning 21-040) in a dose range of (6.25–50 μg/mL). Digitonin solutions were incubated on fixed neurons for 10 minutes at room temperature to induce plasma membrane permeabilization. Following digitonin treatment, neurons were incubated with 2% bovine serum albumin (Sigma Aldrich A7906) in PBS (Corning 21-040) for 60 minutes for blocking. A full permeabilization condition was conducted in parallel in accordance with the general ICC protocol. Thus, all wells were blocked in 2% BSA for a total of 60 minutes. Digitonin wells were incubated with anti-TGN46, anti-GM130, and anti-MAP2 antibodies in a 2% BSA solution in PBS at dilutions described in **Table S1** overnight at 4 °C. Control wells were incubated with the same antibodies and dilutions overnight at 4 °C but in blocking buffer described in the general protocol. The following day all wells were washed as described in the general protocol. Both the control and experimental wells received secondary antibodies and stains at the concentrations listed in **Table S2**, with digitonin secondary solution being diluted in 2% BSA in PBS. All remaining steps were carried out in accordance with the general protocol. Confocal fluorescence microscopy images were obtained using a Yokogawa CellVoyager CQ1 spinning disk confocal microscope. All images shown represent the maximum projection from a Z-stack imaging technique. Laser power and exposure times were identical within a set of experiments.

For the topology experiment: All steps were identical to the optimization experiment except that only the optimal digitonin concentration (25 μg/mL) was used for the partial permeabilization condition, and anti-5-HT_2A_ R and anti-INMT antibodies were used instead of anti-TGN46, as this enzyme served as a topology marker. All other steps were identical.

### Bulk spatial proximity analysis per image (Figure 2b)

Images were analyzed in Fiji using the BIOP-JACoP plugin under default settings, and each spatial proximity analysis description generated reflects results across an entire image.

### Image analysis using an object-based spatial proximity analysis pipeline (Figure 3c-e & Figure 6d,e)

To quantify spatial proximity, images were analyzed using Fiji software with our object-based ROI generation (**Code S1**). ROIs encapsulating whole neurons were generated by combining MAP2 and Hoechst-derived binary masks. Exact threshold cutoffs differed between experiments to account for different imaging conditions and plate variability (**Table S3**). As a preprocessing step, certain channels were subject to an intensity cap to deflate the influence of the top 1-2% of pixels that were artifacts of the ICC protocol (**Table S3**). After generating neuron ROIs and preprocessing, the BIOP-JACoP plugin was used to generate the Pearson’s and Manders’ correlation coefficients for each ROI. For each antibody the autothresholding operation (Li, Moments, Otsu, or Huang) was chosen based on signal phenotype and histogram distribution and was held constant across all images of that antibody (**Table S3**). To prevent over representation of non-neuronal cell types, all ROIs used were compared to the original MAP2 image to validate neuronal morphology, without viewing other antibody channels to prevent bias.

### Line ROI analysis for *cis*-Golgi localization (Figure 4b,c)

Line ROI analysis was performed in Fiji as previously described by van Bommel et al.^42^ Briefly, neurons with Golgi stacks aligned in the imaging plane were selected using only the GM130 channel for visualization to prevent bias, and linear ROIs were drawn perpendicular to the Golgi stack. Immunofluorescence intensity was plotted for GM130 and INMT along the same ROI, baseline-corrected, and peak heights normalized. The distance between the maximum GM130 and INMT intensity peaks was then calculated for each neuron.

### Image analysis for Golgi ROI intensity readouts (Figure 5c)

the image corresponding to GM130 signal was opened in Fiji and duplicated. Using the threshold tool, a pixel intensity range was set manually on the duplicated image to highlight individual Golgi structures, creating a binary mask. Using the set measurements module, area and mean intensity were activated, and the reading was redirected to the channel corresponding to INMT immunoreactivity. Analyze particles was used to generate GM130 ROIs. ROIs corresponding to Golgi structures identified in the original image were recorded for their mean INMT intensity. This process was repeated redirecting the analyze particles to the GM130 channel, and the same ROIs were selected and recorded for their mean GM130 intensity. INMT intensity was normalized to GM130 intensity before plotting.

### Digitonin-based permeabilization experiments image analysis (Figure 7c,d & Figure S7b)

For the optimization experiment: The optimization portion of the workflow aimed at determining the optimal digitonin concentration, so only Golgi structures were analyzed because GM130 and TGN46 have defined topology at the Golgi apparatus. The GM130 channel was used to generate Golgi ROIs through percentile-based thresholding (**Code S2**). The top 1% brightest GM130 pixels were selected based on the intensity histogram and converted into a binary mask. Analyze particles was used to generate discrete Golgi ROIs. Mean signal within each Golgi ROI was assessed for TGN46 intensity using the redirect function in Set Measurements, and values were normalized to the full permeabilization condition to assess membrane accessibility across digitonin concentrations. This analysis was used to determine the optimal digitonin concentration to be 25 μg/mL.

The GM130 and Hoechst channels were used to generate Golgi and nucleus ROIs through percentile-based thresholding (**Code S2**). The top 1% brightest GM130 pixels and top 9% brightest Hoechst pixels were selected based on the intensity histogram and converted into a binary mask. Analyze particles was used to generate discrete Golgi or nucleus ROIs. Mean signal intensity for INMT and 5-HT_2A_ R immunoreactivity was recorded within these ROIs and plotted relative to the full permeabilization condition. Trends in average intensities of INMT and 5-HT_2A_ R immunoreactivity under the partial and full permeabilization conditions were used to determine topology.

### RNA sequencing

Transcriptomic profiling was performed by Metware Biotechnology (Woburn, MA) using an RNA-seq workflow. Total RNA was isolated from WT HEK and HEK-INMT-OE cells with three biological replicates analyzed per condition. RNA extraction, library preparation, and sequencing. The cells were thawed on ice. RNA was extracted using RNeasy Plus Mini kit (Qiagen) following the manufacturer’s recommendations. The quality of isolated RNA was assessed using RNA Tapestation (Agilent Technologies G2991BA) and quantified using Infinite F Nano+ 200 Pro (Tecan). Paramagnetic beads coupled with oligo d(T) are combined with total RNA to isolate poly(A)+ transcripts using NEBNext Poly(A) mRNA Magnetic Isolation Module (New England BioLabs E7490). Prior to first strand synthesis, samples are randomly primed using N6 nucleotides and fragmented based on manufacturer’s recommendations. The first strand is synthesized with the Protoscript II Reverse Transcriptase with extension period set to 30 minutes at 42C. All remaining steps for library construction were performed according to the NEBNext Ultra II Directional RNA Library Prep Kit for Illumina (New England BioLabs E7760). Final library was quantified using Qubit 2.0 (ThermoFisher Q32866) and the library quality was assessed by TapeStation (Agilent Technologies G2991BA). Equimolar pooling of libraries was performed based on quantification values and sequenced on an Illumina NovaSeq X plus (Illumina) with a read length configuration of 150 PE for 20 million PE reads per sample. Raw sequencing reads were filtered using fastp to remove adapter sequences, low-quality reads, and reads containing >10% ambiguous bases. Clean reads were aligned to the human reference genome (Homo sapiens GRCh38) using HISAT2, and genelevel read counts were quantified using featureCounts. Gene expression levels were reported as fragments per kilobase of transcript per million mapped reads (FPKM). Differential gene expression analysis between experimental groups was performed using DESeq2, and p-values were adjusted using the Benjamini– Hochberg method to control the false discovery rate (FDR). Genes with |log₂ fold change| ≥ 1 and FDR < 0.05 were considered significantly differentially expressed.

## ABBREVIATIONS

INMT: Indolethylamine *N*-methyltransferase
DMT: *N,N*-dimethyltryptamine
5-HT2A R: Serotonin 2A receptor
CNS: Central nervous system
GPCR: G protein-coupled receptor
NMT: *N*-Methyltryptamine
AADC: Aromatic L-amino acid decarboxylase
MAO-A: Monoamine oxidase A
mRNA: Messenger ribonucleic acid
snRNA-seq: Single cell RNA sequencing
ISH: *In situ* hybridization
hINMT: Human INMT
SAM: *S*-adenosyl-L-methionine
KO: Knockout
SAH: *S*-adenosyl-L-homocysteine
iPSC: Induced pluripotent stem cell
ICC: Immunocytochemistry
WT HEK: Wildtype HEK293T cell line
HEK-INMT-OE: INMT-overexpressing HEK293T cell line
BIOP-JACoP: BioImaging and Optics Platform Just Another Colocalization Plugin
kDa: Kilodalton
GAPDH: Glyceraldehyde 3-phosphate dehydrogenase
shRNA: Short hairpin RNA
GFP: Green fluorescent protein
MAP2: Microtubule-associated protein
2 ROI: Region of interest
GM130: Golgi matrix protein 130
ER: Endoplasmic reticulum
TGN46: Trans-Golgi network protein 46

## AUTHOR INFORMATION

### Author Contributions

K.W. and S.J.H. conceived of the project. K.W. performed computational analysis. K.W. performed cell maintenance and immunocytochemistry experiments, with design input from S.J.H. and S.A.R. K.W. performed viral overexpression and knockdown experiments. S.L.M. performed immunoblot assays. K.W. designed and performed permeabilization assays. K.W. designed image analysis pipelines for all immunocytochemistry experiments with input from S.A.R. K.W. analyzed immunocytochemistry data. K.W. assembled figures. K.W. wrote manuscript text with input from S.L.M., S.A.R., and S.J.H. S.J.H. and K.W. held overall responsibility for the study.

## ACKNOWLEDGEMENT

We thank Professor Brian Liau and Professor David Liu for their scientific advice on this project. We thank Professor Martin Kuchař for generously providing the anti-DMT-like antibody.

