## Supplemental Information for "Mapping Endogenous Psychedelic Chemistry: Indolethylamine *N*-Methyltransferase (INMT) Localizes Proximal to the Golgi Apparatus of Human Neurons"

**AUTHOR INFORMATION**

Karis Will[1,2,3], Sergio Lázaro Martínez[1,2,4], Surya A. Reis[1,2], Stephen J. Haggarty[1,2*]

**AFFILIATIONS**

1. Chemical Neurobiology Laboratory, Center for Genomic Medicine, Department of Neurology, Massachusetts General Hospital, Harvard Medical School, Boston, MA, USA.

2. Center for the Neuroscience of Psychedelics, Department of Psychiatry, Massachusetts General Hospital, Boston, MA, USA.

3. Department of Chemistry and Chemical Biology, Harvard University, Cambridge, MA, USA.

4. Department of Psychiatry & Neuropsychology, School for Mental Health and Neuroscience, Maastricht University, Maastricht, Limburg, The Netherlands.

*Correspondence should be addressed to:

Stephen J. Haggarty, Professor of Neurology

Massachusetts General Hospital, Harvard Medical School

Center for Genomic Medicine

185 Cambridge Street, Boston, MA, 02114, USA

**Supplemental Material**
(21 pages)

***Contents***

**Schematic S1** - Proposed reaction mechanism of hINMT methylation of tryptamine
**Figure S1** - Immunogen competition assay in TT cells
**Figure S2** - Immunogen competition assay in human iPSC-derived neurons
**Figure S3** - pLV-Puro-EF1α-hINMT-FLAG vector map and hINMT-FLAG coding sequence
**Figure S4** - Complete, unedited immunoblot membranes used to generate **Figure 2c-f**
**Figure S5** - Differentially expressed genes between WT HEK and HEK-INMT-OE cells
**Figure S6** - shRNA-mediated INMT knockdown in HEK293T cells
**Figure S7** - Digitonin concentration optimization
**Table S1** - Primary antibodies and dilutions
**Table S2** - Secondary antibodies and dilutions
**Table S3** - Colocalization image analysis parameters
**Code S1** - Object-based colocalization analysis pipeline
**Code S2** - Object-based mean intensity analysis pipeline

### SUPPLEMENTAL FIGURES

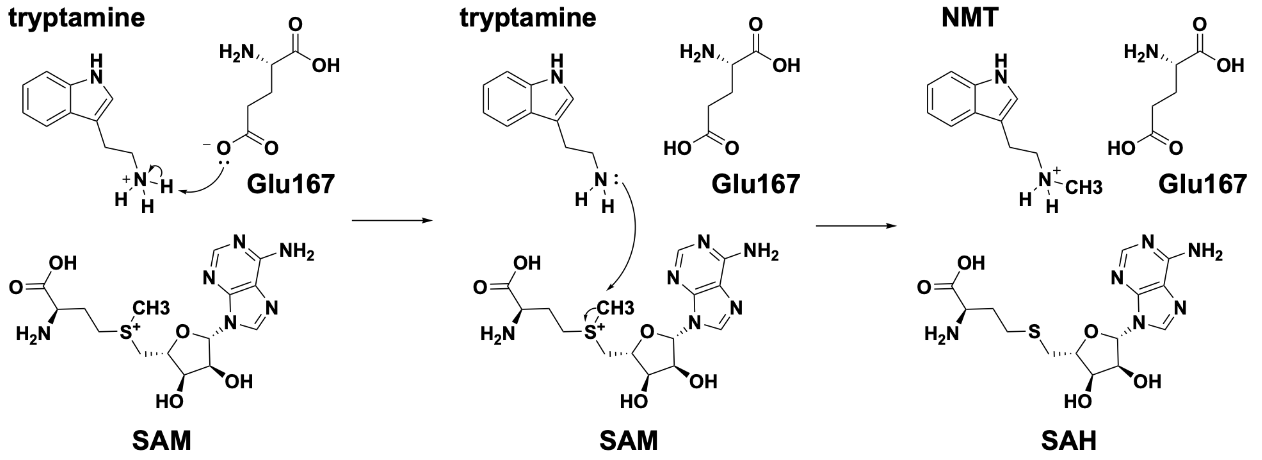


**Schematic S1:** Proposed reaction mechanism of hINMT methylation of tryptamine. SAM - *S*-adenosyl-L-methionine; SAH - *S*-adenosyl-L-homocysteine

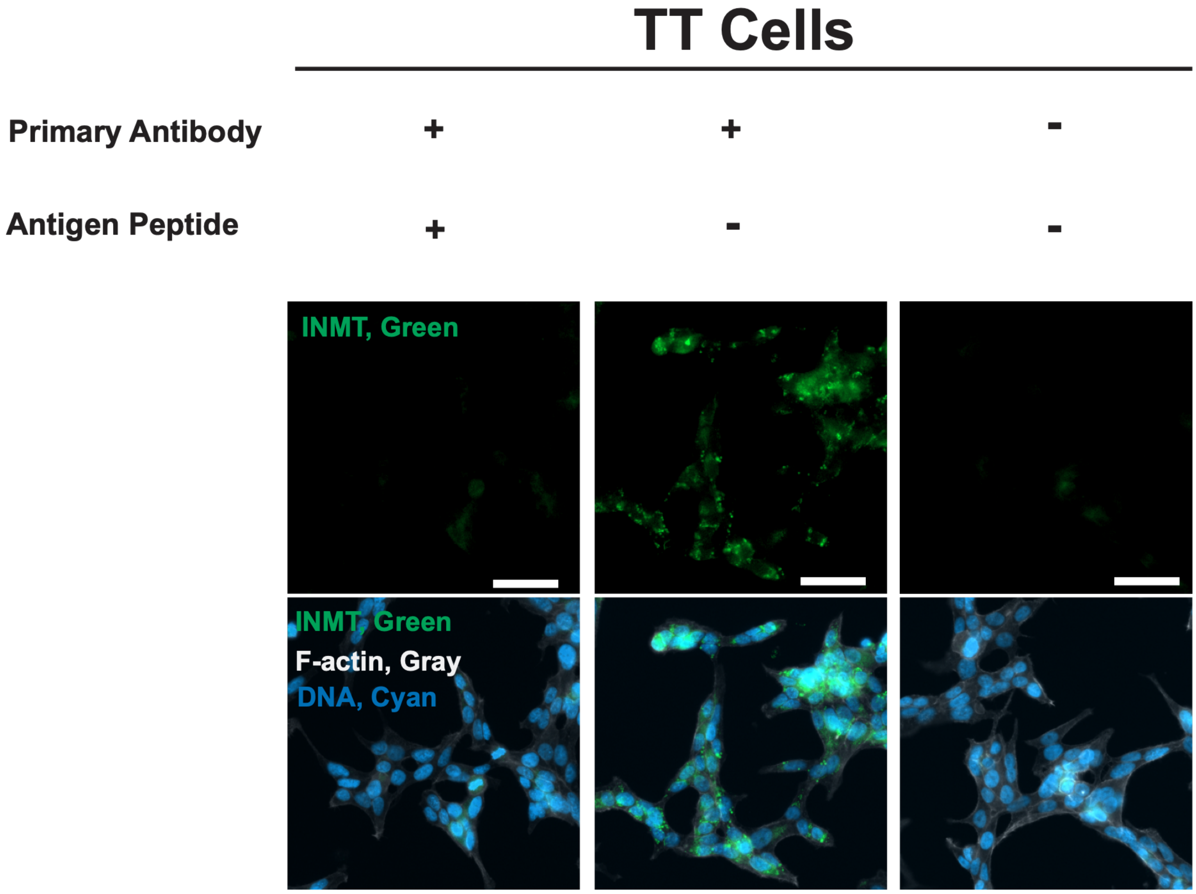


**Figure S1.** Pre-incubation of the anti-INMT antibody with its immunizing peptide reduces immunoreactivity in TT cells**.**

Representative images of TT cells (ATCC CRL-1803) immunolabeled using an anti-INMT antibody. *Left column:* Antibody pre-incubated with its immunizing peptide. *Middle column:* Free antibody without peptide pre-incubation. *Right column:* Secondary antibody only control. Scale bars: 40 μm.

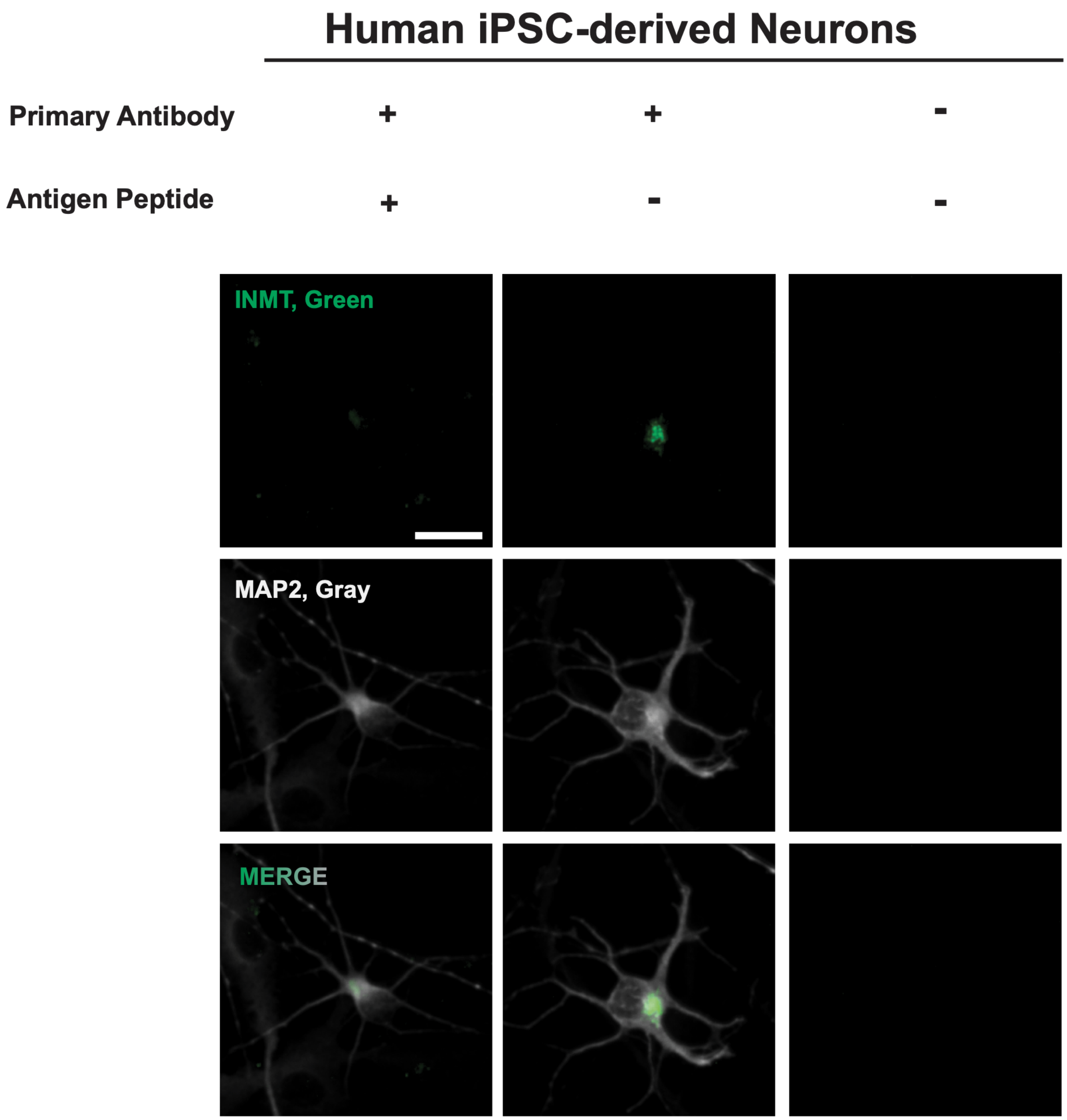


**Figure S2.** Pre-incubation of the anti-INMT antibody with its immunizing peptide reduces immunoreactivity in human iPSC-derived neurons.

Representative images of human iPSC-derived neurons immunolabeled using an anti-INMT antibody. *Left column:* Antibody pre-incubated with its immunizing peptide. *Middle column:* Free antibody without pre-incubation. *Right column:* Secondary antibody only control. Scale bar represents 20 μm.

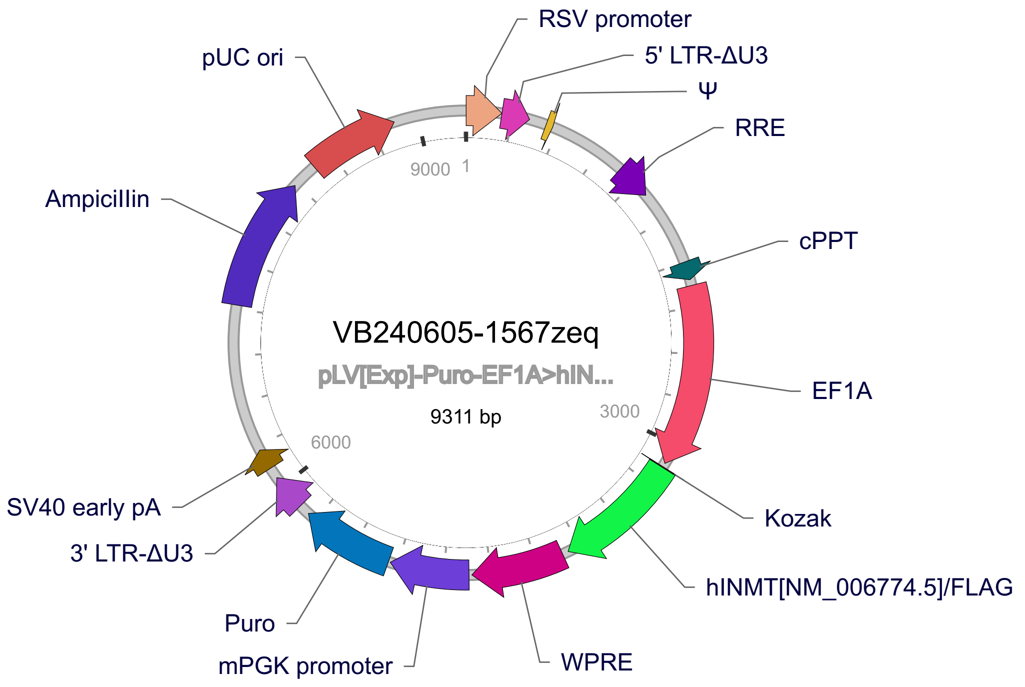


ATGAAGGGTGGCTTCACTGGGGGTGATGAGTACCAGAAGCACTTCCTGCCCAGGG
ACTACTTGGCTACTTACTACAGCTTCGATGGCAGCCCCTCACCCGAGGCCGAGATG
CTGAAGTTTAACTTGGAATGTCTCCACAAGACCTTCGGCCCTGGAGGCCTCCAAGG
GGACACGCTGATTGACATTGGCTCAGGTCCTACCATCTACCAAGTTCTTGCTGCCT
GTGATTCCTTCCAAGACATCACTCTCTCCGACTTTACCGACCGCAACCGGGAGGAG
CTGGAAAAGTGGCTGAAGAAGGAGCCGGGGGCCTATGACTGGACCCCAGCGGTG
AAATTCGCCTGTGAGCTGGAAGGAAACAGCGGCCGATGGGAGGAGAAGGAGGAG
AAGCTGCGGGCAGCGGTGAAGCGGGTGCTCAAGTGCGATGTCCACCTGGGCAAC
CCGCTGGCCCCGGCTGTGTTGCCTCTCGCCGACTGTGTGCTCACCCTGCTGGCCA
TGGAGTGTGCCTGCTGTAGCCTTGATGCCTACCGCGCTGCCCTGTGCAACCTTGC
CTCACTGCTCAAGCCGGGTGGCCACCTGGTGACCACTGTCACGCTTCGGCTCCCG
TCCTACATGGTGGGGAAGCGTGAATTTTCCTGCGTGGCCCTGGAGAAAGAGGAGG
TGGAGCAGGCTGTCCTGGATGCTGGCTTTGACATTGAACAGCTCCTACACAGTCCC
CAGAGCTACTCTGTCACCAATGCTGCCAACAATGGGGTCTGCTTCATTGTGGCTCG
CAAGAAGCCTGGGCCCGACTACAAAGACGATGACGACAAGTGA

**Figure S3.** pLV-Puro-EF1α-hINMT-FLAG vector map and hINMT-FLAG coding sequence.

**a** **b**

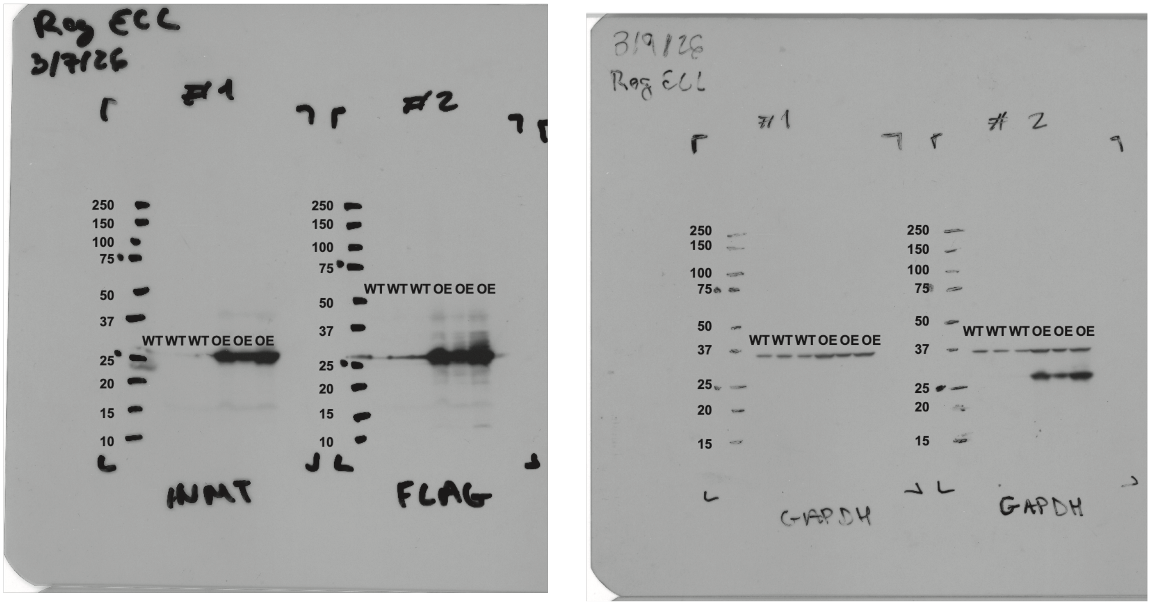


**Figure S4:** Complete, unedited membranes of cell lysate samples collected from WT HEK (WT) and HEK-INMT-OE (OE) cells using **a)** anti-INMT (left) and anti-FLAG (right) or **b)** anti-GAPDH antibodies.

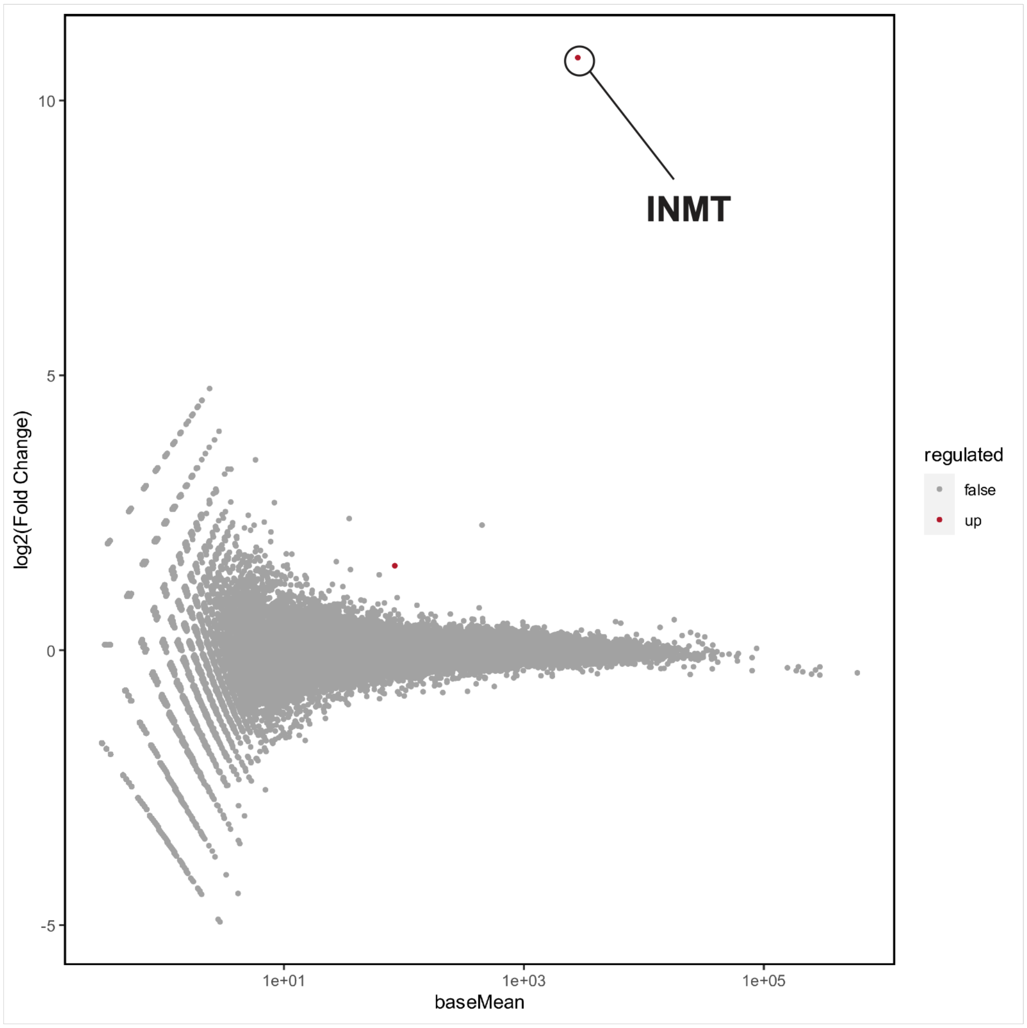


**Figure S5**: MA (mean-average) plot of differentially expressed genes between WT HEK and HEK-INMT-OE cell lines.

The horizontal coordinate indicates the mean value of gene expression in the two samples; red dots represent up-regulated gene expression, while blue dots represent down-regulated gene expression, and gray indicates no significant difference in gene expression.

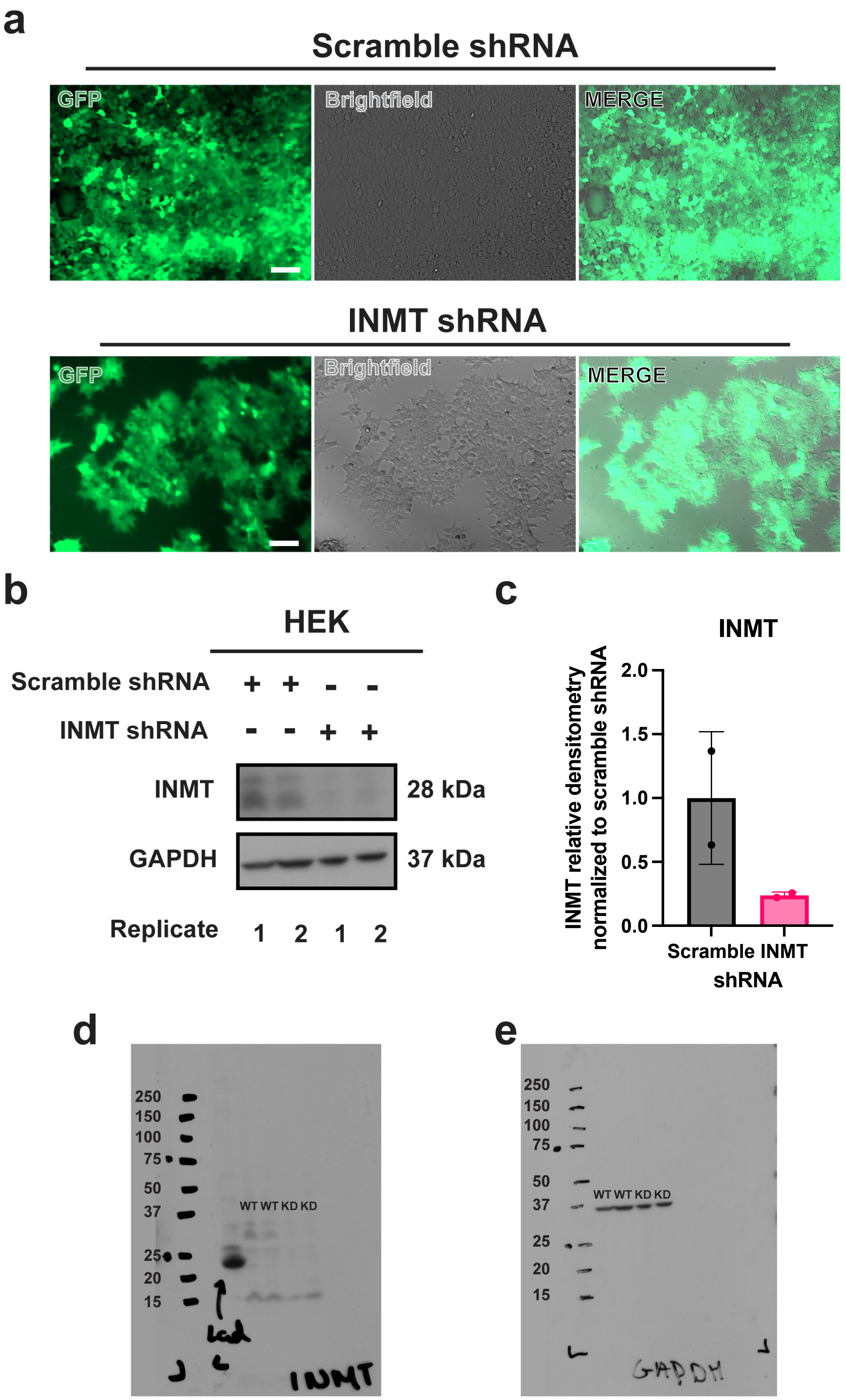


**Figure S6:** shRNA-mediated INMT knockdown in HEK293T cells supports antibody ability to bind to INMT in human cellular contexts.

**a)** Representative fluorescence and brightfield images of HEK293T cells transfected with control scramble shRNA (top) or INMT targeting shRNA (bottom). GFP channel reflects degree of transduction. **b)** Immunoblot illustrating INMT immunoreactivity in HEK293T cells transduced with scramble or INMT targeting shRNA. **c)** Densitometric quantification of INMT immunoreactivity, relative to GAPDH, in HEK293T cells transduced with scramble or INMT targeting shRNA. Densitometric quantification was normalized to the scramble shRNA condition. Bars represent mean ± SD; dots represent *n* = 2 biological replicates. Complete, unedited membranes of cell lysate samples collected from HEK293T cells transduced with scramble shRNA (WT) or INMT targeting shRNA (KD) using **d)** anti-INMT or **e)** anti-GAPDH antibodies. Membranes were developed using SuperSignal West Pico Extended Duration Substrate. GAPDH - Glyceraldehyde 3-phosphate dehydrogenase.

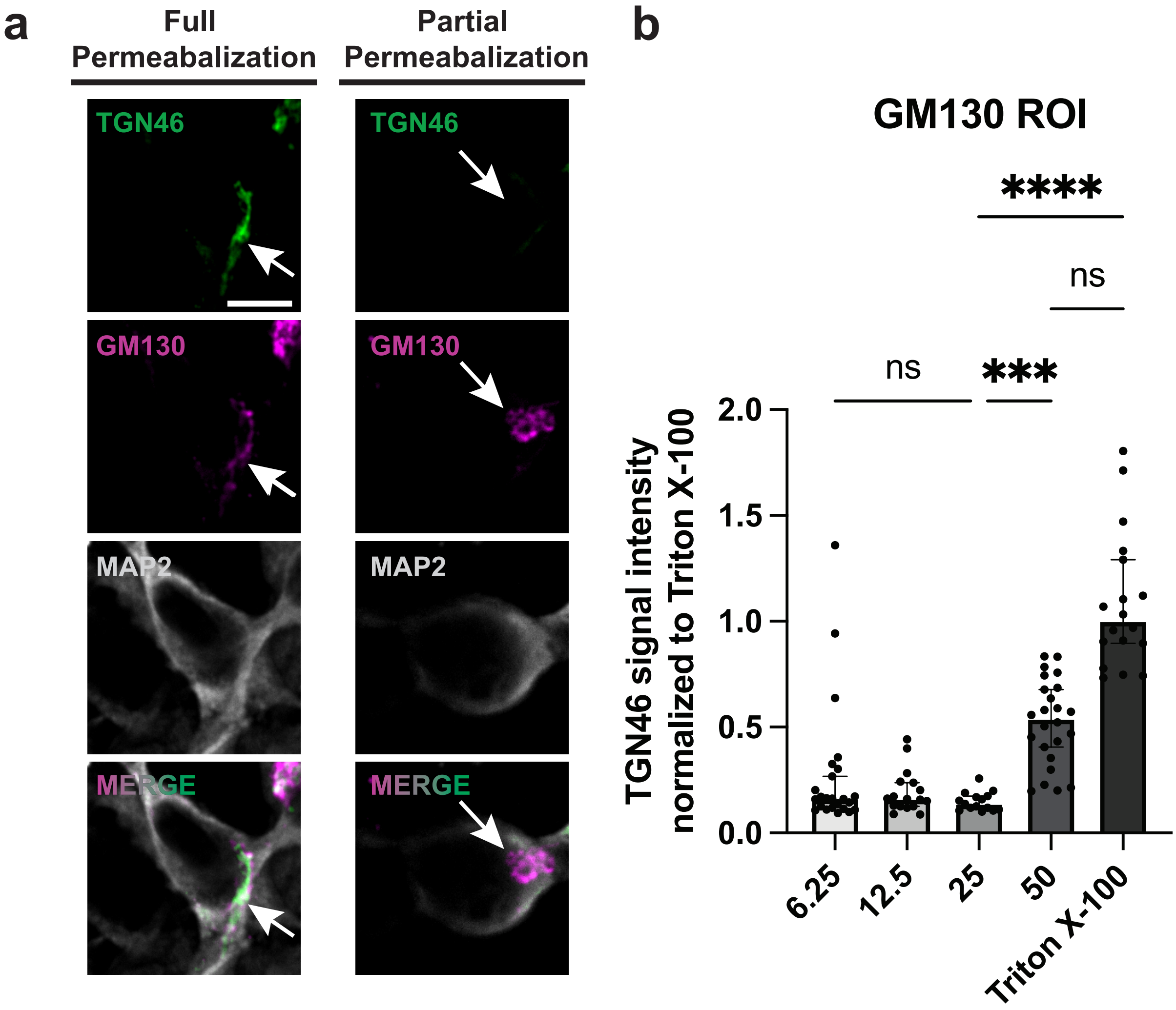


**Figure S7:** Digitonin concentration optimization.

Representative images of human iPSC-derived neurons permeabilized with either **a)** 0.3% Triton X-100 or 25 µg/mL digitonin and co-incubated with anti-TGN46, anti-GM130, and anti-MAP2 antibodies. **b)** Quantification of mean TGN46 intensities plotted as a function of digitonin concentration, normalized to the Triton X-100 full permeabilization condition. Bars show median ± 95% CI; dots represent mean immunoreactivity from a single ROI. ns = not significant; ***p < 0.001; ****p < 0.0001; Kruskal–Wallis tests followed by Dunn's multiple comparisons tests; *n* = 15 - 24 cells per condition from one independent experiment.

### SUPPLEMENTAL METHODS

**Table S1:** Primary antibodies and dilutions

| Antibody | Host | Vendor | Catalog # | Application (dilution) |
|---|---|---|---|---|
| anti-INMT | Rabbit | Novus Biologicals | NBP2-57073 | ICC (1:100); WB (1:500) |
| anti-AADC | Mouse | Invitrogen | MA5-31379 | ICC (1:100) |
| anti-MAO-A | Mouse | Santa Cruz | sc-271123 | ICC (1:50) |
| anti-MAP2 | Chicken | EnCor Biotechnology | CPCA-MAP2 | ICC (1:1000) |
| anti-GM130 | Mouse | Abcam | ab169276 | ICC (1:50) |
| anti-TGN46 | Sheep | Bio-Rad | AHP500G | ICC (1:250) |
| anti-5-$HT_{2A}$ R | Rabbit | Neuromics | RA24288 | ICC (1:50) |
| anti-DMT | Rabbit | Gift (Martin Kuchař) | — | ICC (1:25) |
| anti-FLAG | Mouse | Sigma-Aldrich | F1804 | ICC (1:75); WB (1:500) |
| anti-GAPDH | Mouse | Abcam | ab8245 | WB (1:10,000) |

**Table S2:** Secondary antibodies and dilutions

| Antibody | Target species | Tag | Vendor | Catalog # | Application (dilution) |
|---|---|---|---|---|---|
| anti-rabbit 488 | Rabbit | Alexa Fluor 488 | Invitrogen | A32731 | ICC (1:500) |
| anti-mouse 647 | Mouse | Alexa Fluor 647 | Invitrogen | A21235 | ICC (1:500) |
| anti-chicken 594 | Chicken | Alexa Fluor 594 | Invitrogen | A11042 | ICC (1:500) |
| anti-sheep 488 | Sheep | Alexa Fluor 488 | Invitrogen | A11015 | ICC (1:500) |
| anti-rabbit HRP | Rabbit | Horseradish peroxidase | Cell Signaling | 7074S | WB (1:4000) |
| anti-mouse HRP | Mouse | Horseradish peroxidase | Cell Signaling | 7076S | WB (1:4000) |

**Table S3:** Colocalization image analysis parameters

| Analysis Pair (a-b) | Trim | DAPI Threshold | MAP2 Threshold | Channel A Autothreshold | Channel B Autothreshold | Figure |
|---|---|---|---|---|---|---|
| INMT - MAOA | 0.99 INMT 1.00 MAOA | 1200 | 1200 | Li | Moments | 3 |
| INMT - AADC | 0.99 INMT 0.99 AADC | 1200 | 1700 | Li | Moments | 3 |
| INMT - GM130 | 0.99 INMT 1.00 GM130 | 2200 | 1600 | Li | Li | 6 |
| INMT - Hoechst | 0.99 INMT 0.98 Hoechst | 2200 | 1600 | Li | Otsu | 6 |
| 5-$HT_{2A}$ R - GM130 | 1.00 5-$HT_{2A}$ R 1.00 GM130 | 2200 | 1600 | Huang | Li | 6 |
| 5-$HT_{2A}$ R - Hoechst | 1.00 5-$HT_{2A}$ R 0.98 Hoechst | 2200 | 1600 | Huang | Otsu | 6 |
| DMT - GM130 | 0.99 DMT 1.00 GM130 | 2200 | 1600 | Moments | Li | 6 |
| DMT - Hoechst | 0.99 DMT 0.98 Hoechst | 2200 | 1600 | Moments | Otsu | 6 |

**Code S1.** Object-based colocalization analysis pipeline.

```
INPUT_DIR  = "InputFolder”;
OUTPUT_DIR = "OutputFolder’;
if (!File.exists(OUTPUT_DIR)) File.makeDirectory(OUTPUT_DIR);

TRIM_C1 = 1.0;
TRIM_C2 = 0.99;
TRIM_C3 = 1.0;
TRIM_C4 = 0.99;
HARD_REMOVE  = false;

COSTES_BLOCK = 5;
```

```
COSTES_SHUFF = 100;

MAP2_FIXED_LO = 1700;
DAPI_FIXED_LO = 1200;

function parseWF(titleStr) {
  dot = indexOf(titleStr, ".tif");
  if (dot < 0) dot = indexOf(titleStr, ".tiff");
  if (dot >= 0) titleStr = substring(titleStr, 0, dot);
  iW = indexOf(titleStr, "W"); if (iW < 0) exit("No 'W' in title: "+titleStr);
  rest = substring(titleStr, iW+1);
  iFrel = indexOf(rest, "F"); if (iFrel < 0) exit("No 'F' in title: "+titleStr);
  wellStr = substring(rest, 0, iFrel);
  rest2 = substring(rest, iFrel+1);
  iTrel = indexOf(rest2, "T"); if (iTrel < 0) exit("No 'T' in title: "+titleStr);
  fldStr = substring(rest2, 0, iTrel);
  return newArray(parseInt(wellStr), parseInt(fldStr), "W"+wellStr, "F"+fldStr);
}

function capTopX(srcTitle, trimVal) {
  if (trimVal >= 1) return srcTitle;
  selectImage(srcTitle);
  run("Duplicate...", "title="+srcTitle+"_cap");
  selectImage(srcTitle+"_cap");
  getHistogram(v, h, 4096);
  tot=0; for(i=0;i<h.length;i++) tot+=h[i];
  target=tot*trimVal;
  cum=0; cut=v[v.length-1];
  for(i=0;i<h.length;i++){ cum+=h[i]; if(cum>=target){ cut=v[i]; break; } }
  setThreshold(cut, 65535, "raw"); run("Create Selection");
  if (HARD_REMOVE) run("Set...", "value=0"); else run("Set...", "value="+cut);
```

```
  run("Select None"); resetThreshold();
  return srcTitle+"_cap";
}

function backfillROIStatsIntoLastRows(pairTag, WELL_ID, FLD_ID) {
  startRow = nResults - nROI;
  if (startRow < 0) startRow = 0;
  for (r=0; r<nROI; r++) {
    row = startRow + r;
    setResult("Area", row, area[r]);
    setResult("Mean", row, mean[r]);
    setResult("Min", row, minv[r]);
    setResult("Max", row, maxv[r]);
    setResult("X", row, cx[r]);
    setResult("Y", row, cy[r]);
    setResult("Perim.", row, perim[r]);
    setResult("Circ.", row, circ[r]);
    setResult("Feret", row, feret[r]);
    setResult("FeretX", row, feretX[r]);
    setResult("FeretY", row, feretY[r]);
    setResult("FeretAngle", row, feretAng[r]);
    setResult("MinFeret", row, minFeret[r]);
    setResult("AR", row, ar[r]);
    setResult("Round", row, roundv[r]);
    setResult("Solidity", row, solidity[r]);
    setResult("Well_ID", row, WELL_ID);
    setResult("Fld_ID", row, FLD_ID);
    setResult("Pair", row, pairTag);
  }
  updateResults();
}
```

```
c1_name = "FileNameC1.tif";
c2_name = "FileNameC2.tif";
c3_name = "FileNameC3.tif";
c4_name = "FileNameC4.tif";

open(INPUT_DIR + c1_name);
open(INPUT_DIR + c2_name);
open(INPUT_DIR + c3_name);
open(INPUT_DIR + c4_name);

wf = parseWF(c3_name);
WELL_ID  = wf[0];
FLD_ID   = wf[1];
WELL_TAG = wf[2];
FLD_TAG  = wf[3];

selectImage(c3_name);
run("Duplicate...", "title=MAP2_tmp");
selectImage("MAP2_tmp");
setOption("BlackBackground", true);
setThreshold(MAP2_FIXED_LO, 65535, "raw");
run("Convert to Mask");
rename("MAP2_mask");

selectImage(c1_name);
run("Duplicate...", "title=DAPI_tmp");
selectImage("DAPI_tmp");
setOption("BlackBackground", true);
setThreshold(DAPI_FIXED_LO, 65535, "raw");
```

```
run("Convert to Mask");
rename("DAPI_mask");

run("Image Calculator...", "operation=OR create image1=MAP2_mask
image2=DAPI_mask");
rename("ROI_mask");
setOption("BlackBackground", true);
run("Convert to Mask"); // ensure binary

run("Set Measurements...", "area mean min centroid perimeter shape feret's
redirect="+c3_name+" decimal=3");
selectImage("ROI_mask");
run("Analyze Particles...", "size=2000-Infinity pixel display clear include add");

nROI = roiManager("count");
for (i=0; i<nROI; i++) { roiManager("select", i); roiManager("Rename", ""+(i+1)); }

run("Clear Results");
selectImage(c3_name);
for (i=0; i<nROI; i++) { roiManager("select", i); run("Measure"); }

area=newArray(nROI); mean=newArray(nROI); minv=newArray(nROI);
maxv=newArray(nROI);
cx=newArray(nROI); cy=newArray(nROI); perim=newArray(nROI);
circ=newArray(nROI);
feret=newArray(nROI); feretX=newArray(nROI); feretY=newArray(nROI);
feretAng=newArray(nROI); minFeret=newArray(nROI); ar=newArray(nROI);
roundv=newArray(nROI); solidity=newArray(nROI);

for (i=0;i<nROI;i++){
  area[i]=getResult("Area",i); mean[i]=getResult("Mean",i); minv[i]=getResult("Min",i);
```

```
  maxv[i]=getResult("Max",i); cx[i]=getResult("X",i); cy[i]=getResult("Y",i);
  perim[i]=getResult("Perim.",i); circ[i]=getResult("Circ.",i); feret[i]=getResult("Feret",i);
  feretX[i]=getResult("FeretX",i); feretY[i]=getResult("FeretY",i);
  feretAng[i]=getResult("FeretAngle",i); minFeret[i]=getResult("MinFeret",i);
  ar[i]=getResult("AR",i); roundv[i]=getResult("Round",i); solidity[i]=getResult("Solidity",i);
}
run("Clear Results");

if (isOpen("MAP2_mask")) close("MAP2_mask");
if (isOpen("DAPI_mask")) close("DAPI_mask");
if (isOpen("ROI_mask"))  close("ROI_mask");
if (isOpen("MAP2_tmp"))  close("MAP2_tmp");
if (isOpen("DAPI_tmp"))  close("DAPI_tmp");

protein1_cap = capTopX(c1_name, TRIM_C1);
protein2_cap = capTopX(c2_name, TRIM_C2);
protein3_cap  = capTopX(c3_name, TRIM_C3);
protein4_cap  = capTopX(c4_name, TRIM_C4);

run("Merge Channels...", "c1="+protein2_cap+" c2="+protein4_cap+" create keep");
selectWindow("Composite"); rename("Protein2_vs_Protein4");
selectImage("Protein1_vs_Protein2");
run("BIOP JACoP",
  "channel_a=1 channel_b=2 threshold_for_channel_a=Li
threshold_for_channel_b=Moments " +
  "manual_threshold_a=0 manual_threshold_b=0 crop_rois get_pearsons get_manders
get_fluorogram " +
  "costes_block_size="+COSTES_BLOCK+"
costes_number_of_shuffling="+COSTES_SHUFF);
backfillROIStatsIntoLastRows("protein2_protein4", WELL_ID, FLD_ID);
close("Protein2_vs_Protein4");
```

```
wellDir = OUTPUT_DIR + WELL_TAG + "/";
if (!File.exists(wellDir)) File.makeDirectory(wellDir);
resultsPath = wellDir + "Results_" + FLD_TAG + ".xls";

saveAs("Results", resultsPath);
print("Saved field results to: " + resultsPath);
```

**Code S2:** Object-based mean intensity analysis pipeline.

GM130:

```
selectImage("GM130_Raw_Image.tif");
percentageCutoff = 0.99;
run("Duplicate...", "title=GM130");
selectWindow("GM130");
run("8-bit");

getRawStatistics(nPixels, mean, min, max, std, histogram);
thresholdCount = nPixels * percentageCutoff;
cumulative = 0;
threshold = 0;
for (i = 0; i < 256; i++) {
   cumulative += histogram[i];
   if (cumulative >= thresholdCount) {
      threshold = i;
      break;
   }
}
setThreshold(threshold + 1, 255);
run("Convert to Mask");
rename("Top2percent_Mask");
```

```
selectImage("Top2percent_Mask");
run("Set Measurements...", "area mean integrated redirect= MAP2_Raw_Image.tif decimal.tif decimal=3");
run("Analyze Particles...", "  show=Outlines display clear include add");
run("Merge Channels...", "c4= POI_Raw_Image.tif c6= GM130_Raw_Image.tif create keep");
```

Hoechst:

```
selectImage("Hoechst_Raw_Image.tif");
percentageCutoff = 0.91;
run("Duplicate...", "title=Hoechst");
selectWindow("Hoechst");
run("8-bit");

getRawStatistics(nPixels, mean, min, max, std, histogram);
thresholdCount = nPixels * percentageCutoff;
cumulative = 0;
threshold = 0;
for (i = 0; i < 256; i++) {
   cumulative += histogram[i];
   if (cumulative >= thresholdCount) {
      threshold = i;
      break;
   }
}
setThreshold(threshold + 1, 255);
run("Convert to Mask");
rename("Top2percent_Mask");
selectImage("Top2percent_Mask");
run("Set Measurements...", "area mean integrated redirect= MAP2_Raw_Image.tif decimal.tif decimal=3");
```

```
run("Analyze Particles...", "  show=Outlines display clear include add");
run("Merge Channels...", "c4= POI_Raw_Image.tif c6=Hoechst_Raw_Image.tif create keep");
```
